# Comparative evaluation of synthetic cytokines for enhancing production and performance of NK92 cell-based therapies

**DOI:** 10.1101/2023.05.13.540611

**Authors:** Simrita Deol, Patrick S. Donahue, Roxanna E. Mitrut, Iva J. Hammitt-Kess, Jihae Ahn, Bin Zhang, Joshua N. Leonard

## Abstract

Autologous immune cell therapies are potentially curative, but cost and manufacturing complexity limit access. Off-the-shelf therapies may address these gaps, but further development is needed. The NK92 cell line has a natural killer-like phenotype, has efficacy in cancer clinical trials, and is safe after irradiation. However, NK92 lose activity post-injection, limiting efficacy. This may be addressed by engineering NK92 to express stimulatory factors, for which comparative analysis is needed. Thus, we systematically explored expression of synthetic cytokines for enhancing NK92 production and performance. All synthetic cytokines evaluated (membrane-bound IL2 and IL15, and engineered versions of Neoleukin-2/15, IL15, IL12, and decoy resistant IL18) enhanced NK92 cytotoxicity. Genetically modified cells preferentially expanded by expressing membrane-bound but not soluble synthetic cytokines, and all engineered cells remained sensitive to irradiation. Interestingly, some membrane-bound cytokines conferred cell-contact independent paracrine activity partly attributable to extracellular vesicles. Finally, we characterized interactions within consortia of differently engineered NK92.

## INTRODUCTION

Cell-based cancer immunotherapies have been transformative for certain patients, establishing the importance of addressing challenges that limit their widespread use. Cell-based cancer immunotherapies comprise immune cells with enhanced capacity to kill tumor cells.^1^ Currently, chimeric antigen receptor (CAR) T cells are the most developed and well-studied cell-based cancer therapies.^2^ CAR-T cells can achieve long term remissions in patients unresponsive to multiple lines of chemotherapy, marking a major advance in cancer therapy.^2^ Widespread use of CAR-T cells is limited by high cost and complexity of production.^3^ CAR-T cells must be genetically identical to the patient to avoid graft-versus-host disease, a complication where grafted T cells attack healthy tissue^4^. Therefore, CAR-T cells are typically produced in an autologous fashion—using the patient’s T cells—which requires between 9-14 days^5^ and involves complex supply chains^6^. While mitigating these challenges is the subject of ongoing development and research, it also highlights opportunities for developing complementary approaches.

Natural killer (NK) cell-based therapies are promising alternatives to T cell-based therapies. They confer many comparable benefits (e.g., ability to kill tumor cells) but, unlike T cells, do not need to be genetically identical to the patient, and therefore have the potential to be an “off the shelf” therapy, derived from either allogeneic donors or immortalized cells.^7^ Like CAR-T cells, CAR-NK cells lyse tumor cells and confer durable remissions in patients.^8^ NK cell-based therapies have the additional benefits of causing only mild side effects^8^ and naturally recognizing markers of tumor cells, such as low expression of major histocompatibility complexes^9^. There are multiple sources for NK cells, including the immortalized NK92 cell line, which is attractive for some applications. NK92 cells, which are derived from a non-Hodgkin lymphoma, can replicate for long periods of time, demonstrate NK cell like behavior, express high levels of cytotoxicity activating receptors, and are relatively easy to engineer compared to primary NK cells.^10,11^ After irradiation to prevent tumorigenesis, NK92 are safe to administer and show therapeutic promise in clinical studies.^12^ Notably, NK92 therapies are substantially cheaper to produce than CAR-T cells (less than $20,000 versus $250,000 or greater per treatment).^10^ Finally, NK92 can be repeatedly genetically modified, enabling bioengineers to deliver transgenic cargo above the 10kb limit conferred by standard (e.g., lentiviral) vectors^13^. The increased capacity for modification allows for safety improvements, efficacy improvements, and implementation of sophisticated genetic programs for conferring customizable cellular functions.

Synthetic cytokine (SC) expression may improve production and efficacy of NK92 cell-based therapies. NK92 expansion and anti-tumor efficacy depend on signaling from cytokines^11^, proteins that coordinate immune responses^14^. SCs are analogs of natural cytokines engineered with desirable properties, including improved immune-activating potency and removal of undesirable paradoxical immunosuppressive effects.^15^ SC expression may facilitate NK92 cell-based therapy biomanufacturing by enabling selection and maintenance of engineered cells in recombinant cytokine free culture, simplifying biomanufacturing protocols and requirements.^16^ SC expression could potentially lead to higher NK92 efficacy after administration, enable NK92 to activate surrounding endogenous immune cells, and eliminate the need for systemic co-administration of cytokines, which are only effective at high doses that can lead to toxic side effects^17,18^. To date, some individual SCs have been shown to be effective in enhancing NK92 production and/or performance^19-22^, but a systematic, comparative analysis across SCs is needed to guide future bioengineering—addressing this gap motivates this investigation.

In this study, we evaluated and compared the impact of expressing six SCs^20,23-27^ on properties relevant to NK92 manufacturing (selection and expansion of engineered cells in cytokine-free media), cytotoxicity, and paracrine activity. We evaluated the impact of SC expression on NK92 expansion and cytotoxicity in hypoxia, a feature of solid tumors^28^ that limits efficacy of NK cell-based therapies^29^. Membrane bound, but not soluble, SCs enable selection of engineered cells in cytokine-free media, and effects upon growth and cytotoxicity varied as a function of cytokine choice and environmental condition. Some SCs enabled paracrine activation of NK92 growth or cytotoxicity, and mixing NK92 lines engineered in different ways yielded useful ensemble behaviors. SCs have differential properties and relative advantages which can be used to inform choice of SC in future development in NK92-cell based therapies.

## RESULTS

### Expression and bioactivity of synthetic cytokines

A panel of soluble synthetic cytokines (sSCs) was selected and epitope tagged, then validated for expression and bioactivity on NK92 (**Fig. 1**). The first sSC—Neoleukin-2/15 (Neo2/15)—is an analog of IL2 that preferentially activates pro-inflammatory cells^24^ and, when produced by NK92, stimulates co-cultured immune cells^21^. The second sSC—engineered IL15 (eIL15)—is a fusion of IL15 and the IL15Rα sushi domain that acts as a super agonist on the IL15 receptor^26^, activates NK92 cytotoxicity^30^, and is being evaluated as a monotherapy in clinical trials (NCT05256381). The third sSC—engineered IL12 (eIL12)—is a fusion of the IL12 p35 and p40 monomers (preventing their association into potentially inhibitory cytokines^31^) which potentiates anti-tumor responses in vivo^19^, and, when expressed by NK92, stimulates co-cultured immune cells^19^. The fourth sSC–-decoy-resistant IL18 (DR18)—is a mutated variant of IL18 that does not bind to an inhibitory protein, enhancing its ability to stimulate anti-tumor responses in vivo and activate NK cells.^25^ The sSC transgenes were codon-optimized for expression in human cells with features chosen to enhance stability and add or remove epitope tags as needed (see Methods).

**Figure 1:**
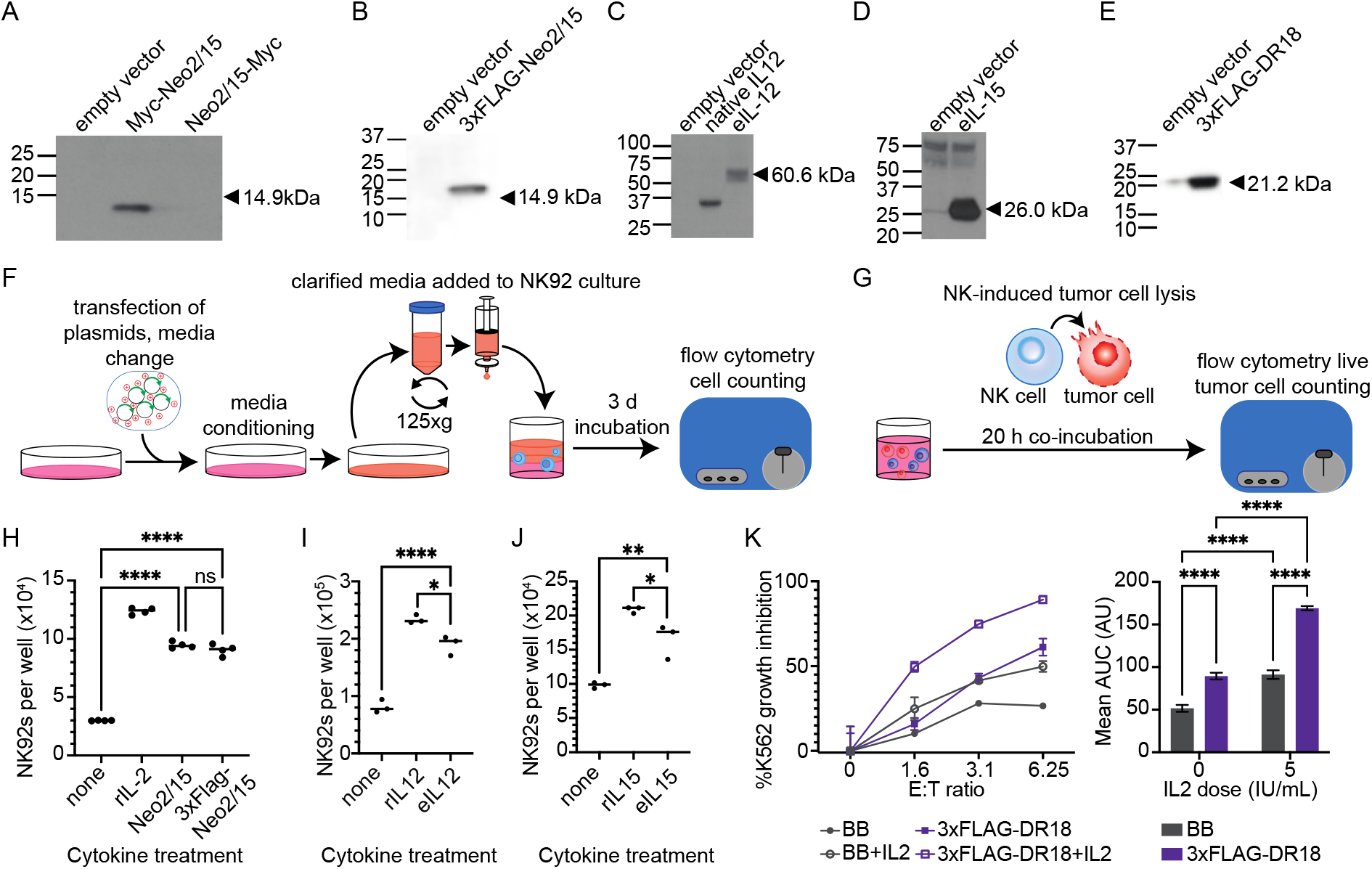
Modified synthetic cytokine constructs are expressed and bioactive. **A-E.** Evaluation of Neo2/15 (Myc tag, **A**), Neo2/15 (3xFLAG tag, **B**), eIL-12 (anti-IL12p40 blot, **C**), eIL-15 (anti IL15 blot, **D**), and DR18 (3XFLAG tag, **E**) expression in HEK293FTs. **F.** Schematic describing the process by which synthetic cytokine bioactivity on NK92 was evaluated. **G.** Schematic describing the assay through which NK92-mediated inhibition of K562 growth (a metric of cytotoxicity) was quantified. **H-J.** Evaluation of bioactivity of synthetic cytokines (Neo2/15, **H**; eIL12, **I**; eIL15, **J**) in HEK293FT-conditioned media. In each case, HEK293FTs were transiently transfected with an empty vector or a vector encoding the indicated synthetic cytokine. Fresh cell medium was conditioned by transfected HEKs for 24 h, starting 48 h after transfection. Conditioned supernatant was harvested and clarified by centrifugation and filtering to remove residual cells and debris. 4×10^4^ NK92 were then cultured in 50% conditioned medium, then counted after 3 d to assess cell growth. Positive controls include NK92 cultured in 50% empty vector conditioned medium supplemented with 100 IU/mL recombinant IL2 (rIL2), 10 ng/mL recombinant IL12 (rIL12), or 10 ng/mL recombinant IL15 (rIL15) as indicated. **** = p<0.0001, ** = p<0.01, ns = p>0.05 (one-way ANOVA, Tukey’s HSD). n=4 (**H**), n=3 (**I**), n=3 (**J**). **K**. Evaluation of bioactivity of DR18 in transduced NK92. NK92 were transduced with an empty backbone lentiviral expression vector encoding no transgene (BB) or that same vector driving expression of 3xFLAG-DR18. NK92 lines were cocultured with K562 cells for 20 h, with and without 5 IU/mL IL2. Live K562 cells counts were measured by flow cytometry and used to calculate %K562 growth inhibition, which is defined by **Equation 1** (see Methods). Points indicate mean values and error bars indicate standard deviation. n=3. Area under the curve (AUC) analysis of each condition was performed. Error bars represent standard error of the mean (SEM) and **** = p<0.0001 (two-way ANOVA, Tukey’s HSD). See **Supplementary Note 1** for full ANOVA results. For all flow-cytometry data, a viability stain was used to exclude dead cells. Abbreviations: arbitrary units (AU), area under the curve (AUC), backbone (BB), international units (IU).

To validate expression of modified sSC constructs by human cells, HEK293FT cells were transiently transfected with sSC constructs and analyzed by western blot. Neo2/15 was only expressed when tagged on its N’ terminus, indicating that Neo2/15 is destabilized when its C terminus is altered (**Fig. 1A**). Neo2/15 was also stably expressed when the N terminal Myc tag was changed to a 3xFLAG tag. 3xFLAG-tagged Neo2/15 (hereafter referred to as Neo2/15) was used for all subsequence experiments (**Fig. 1B**). The eIL12, eIL15, and N’ terminus-tagged DR18 constructs were all expressed by HEK293FTs (**Fig. 1C-E**). Viable constructs were carried forward for further validation.

After validating that modified sSCs were expressed, they were tested for bioactivity on NK92 cells. The native analogs of Neo2/15 (IL2 and IL15), eIL15 (IL15), and eIL12 (IL12) are sufficient to induce NK92 proliferation in the absence of other cytokines^11,32,33^, so it was hypothesized that these sSCs, if bioactive, would also be sufficient to induce NK92 proliferation. To test this property, sSC-conditioned medium was evaluated for its ability to induce NK92 growth in the absence of recombinant IL2, which is usually required for NK92 proliferation (**Fig. 1F**). All three sSCs induced growth, indicating bioactivity (**Fig. 1H-J**). As the native analog of DR18 (IL18) enhances NK92 cytotoxicity in the presence of small amounts of IL2^34^, DR18 should do the same if bioactive. NK92 transduced to express 3xFLAG-tagged DR18 (hereafter referred to as DR18) were tested for their ability to reduce K562 B cell lymphoma cell counts in co-culture (**Fig. 1G**); throughout this study, we use a rigorous metric for quantifying cytotoxicity by comparing the number of live K562 remaining after co-incubation with NK92 to the number of live K562 that would be present if no NK92 were added (**Equation 1**, Methods). DR18 expression significantly enhanced NK92-mediated K562 growth inhibition, particularly in the presence of 5 IU/mL IL2, indicating that DR18 induces autocrine and paracrine bioactivity on NK92 (**Fig. 1K**). Altogether these results indicate that our constructs were functional and fit for subsequent evaluation.

### Selective effects conferred by synthetic cytokine expression

We next investigated whether SC expression enables selection of genetically engineered NK92. We evaluated the previously validated sSCs, as well as two membrane-bound synthetic cytokines (mbSCs). The first mbSC—membrane-bound IL2 (mbIL2)^20^—comprises a fusion of IL2 to IL2Rβ and enhances NK92 cytotoxicity and survival. The second mbSC—membrane-bound IL15 (mbIL15)^27^—is a fusion of IL15 to IL15Rα, enhances NK92 proliferation in vitro^22^ and is related to a membrane-bound IL15 construct that enhanced the proliferation and in vivo cytotoxicity of NK92^35^. Genetic engineering techniques often insert transgenes into a limited fraction of cells, especially as the transgenic cargo increases in size^36^, which can necessitate positive selection (enrichment of genetically engineered cells) to develop an engineered cell product. In contrast, negative selection (depletion of genetically engineered cells or selective expansion of cells that have lack the transduced genes) is the undesired outcome that would pose challenges to developing an engineered cell product. As IL2 and IL15 confer survival of NK92 in the absence of IL2^33^ while IL18 and IL12, despite activating NK92 cytotoxicity^34,37^, also induce NK92 apoptosis when present at higher concentrations^38,39^, we hypothesized that only synthetic counterparts of IL2 or IL15 would impart positive selective pressures on NK92, while DR18 and eIL12 would exert negative selective pressure.

We first evaluated whether expression of SCs would exert selective pressure when starting from a population with a low percentage of transduced cells. NK92 were transduced with lentiviral vectors encoding SCs, cultured in cytokine-free media for 8 d, then evaluated for changes in the percentage of engineered cells (**Fig. 2A**). As Neo2/15 and DR18 lack known signal peptides, which usually direct cytokine trafficking to cellular compartments in which they can bind cytokine receptors^40^, we evaluated Neo2/15 and DR18 constructs with or without an additional N’ terminal IgE signal peptide (notated as sp^IgE^ versus sp^null^, respectively). Since, in preliminary work (not shown), we observed that eIL12 expression downstream of the EF1α promoter induced death of transduced NK92 cells, we evaluated several levels of eIL12 expression using various upstream open reading frames^41^, which led to development of one line capable of sustained expression of eIL12 when cultured with rIL2 (**Supplementary Fig. 1**). After 8 d of culture in cytokine-free media (i.e., without supplementation with recombinant cytokines), only mbIL2 and mbIL15 expression conferred strong positive selection of transduced cells (as compared to populations cultured with rIL2) (**Fig. 2B**). Expression of soluble cytokines did not serve as effective selectable markers, although expression of sp^IgE^Neo2/15 and eIL15 led to slight enrichment of transduced cells (**Fig. 2C**). After 6 weeks of culture without rIL2, the mbIL15 selection conferred greater purity (i.e., frequency of transduced cells in the final population) and higher transgene expression compared to mbIL2 (**Fig. 2D-E**). This outcome may reflect different magnitudes of survival signaling conferred by these mbSCs. A lack of selective advantage was expected for DR18 and eIL12, as neither cytokine is expected to enable long term NK92 survival. A lack of selective advantage for Neo2/15 and eIL15 may be explained by paracrine expansion of non-transduced cells, potentially including competition between transduced and non-transduced cells for a common pool of secreted factors.

**Figure 2:**
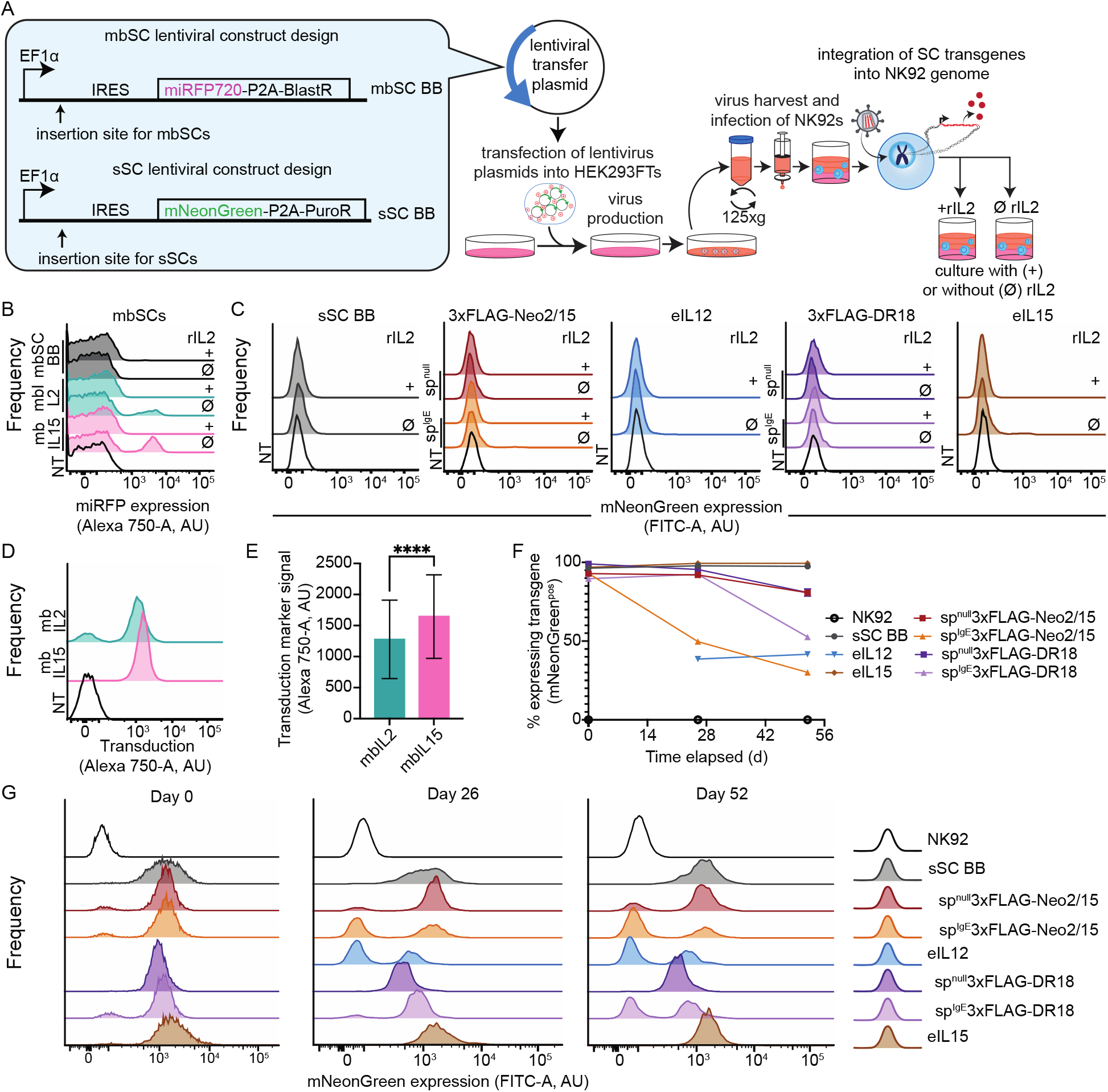
Expression of membrane-bound but not soluble synthetic cytokines confers positive selection on NK92. **A.** Schematic describing NK92 engineering and subsequent selection. **B-C.** Evaluating the potential of SC expression to confer positive selection for mbSCs (**B**) and sSCs (**C**). NK92 transduced to express synthetic cytokines were cultured with or without 100 IU/mL rIL2 for 8 d. Transduced cells were identified by expression of relevant markers (miRFP720 for mbSCs, mNeonGreen for sSCs). **D.** Transgene expression frequency distribution for NK92 expressing mbSC after six weeks of culture without IL2. **E.** Quantification of transgene expression by mbIL2 and mbIL15 NK92 in (**D**). Cells positive for miRFP720^pos^ were gated on and magnitude of miRFP720 signal was analyzed. Bar height indicates mean signal. Error bars represent standard deviation. **** indicates p < 0.0001 (two-tailed unpaired t-test). Cells were analyzed from one sample, each. **F-G.** Evaluating negative selection. Transgene expression was monitored for 52 d. Since cells were sorted on different days but analyzed together, each time point represents a range of days post sorting (see **Supplementary Note 2** for details). As described in the text, eIL12-expressing lines recovered slowly and were not analyzed at day 0. **F** is an analysis of the histograms in **G**. Abbreviations: arbitrary units (AU); backbone (BB), a blasticidin antibiotic resistance gene (BlastR), the constitutive elongation factor 1 α promoter (EF1α), internal ribosome entry site from encephalomycocarditis virus (IRES), membrane-bound synthetic cytokine (mbSC), a near infrared fluorescent protein (miRFP720), a green fluorescent protein (mNeonGreen), non-transduced (NT), puromycin antibiotic resistance gene (PuroR), a 2A self-cleaving peptide from porcine teschovirus-1 polyprotein (P2A), soluble synthetic cytokine (sSC), cells transduced with the indicated sSC lacking a signal peptide (sp^null^); cells transduced with the indicated sSC including an IgE signal peptide (sp^IgE^).

We next evaluated whether expression of sSCs exerted negative selective pressure when starting from a population enriched for transduced cells, as manifested by silencing of the sSC transgenes and expansion of this undesirable population. NK92 engineered to express these factors were enriched by fluorescence activated cell sorting (FACS), and subsequent changes in proportion of engineered cells were tracked over multiple months. eIL15 did not impart negative selective pressures on engineered NK92 but Neo2/15 and DR18 did, particularly when cytokine secretion was enhanced with the IgE signal peptide (**Fig. 2F-G**). Possible explanations include that the latter sSCs may be toxic to NK92 when present above a certain threshold, they may preferentially enhance the expansion of co-cultured cells, or a combination of these effects may exist. We could not similarly evaluate eIL12-expressing NK92 at the earliest time point (“Day 0”; 10 d after FACS enrichment) because cells were present at low number and were slower to recover than other lines, although a stable, low transgene expression frequency was eventually reached (∼40%). This pattern may reflect the toxicity of the eIL12 transgene and its effects after stressors such as FACS. In aggregate, these observations indicate that some sSCs can confer positive selection, some are neutral, and others can be detrimental.

### Effects of membrane-bound synthetic cytokines on NK92 challenged with hypoxia and irradiation

In practice, NK92-based therapies will face challenges that could modulate the effects conferred by synthetic cytokines. In particular, NK92 must be irradiated prior to administration to prevent tumorigenesis^12^, and the hypoxic tumor microenvironment may modulate NK92 cytotoxicity and survival^29^. We thus investigated how signaling from membrane-bound synthetic cytokine (mbSC) expression impacted NK92 growth and when treated with 10 Gy of irradiation (a clinically relevant dose^12^), or under hypoxia.

We first examined the effects of mbSC expression on NK92 expansion in the absence of exogenous cytokine. Both mbIL2 and mbIL15 induced formation of large NK92 clumps, a sign of cell health and proliferation. The mbIL15 NK92 began showing signs of overgrowth (clump spreading and degradation) sooner than did mbIL2 NK92, indicating that mbIL15 NK92 either grew more quickly or were more sensitive to overgrowth (**Fig. 3A, Supplementary Videos 1-3**). Expression of either mbSC increased the growth of NK92 in normoxia (21% O_2_), with mbIL15 trending marginally higher than mbIL2, and marginally improved persistence in hypoxia (1% O_2_) (**Fig. 3B**). Neither mbSC enhanced cell persistence after irradiation with 10 Gy (**Fig. 3B**), indicating that mbIL2 and mbIL15 are not sufficient to confer NK92 resistance to irradiation. This contrasts with prior findings that mbIL2 slightly enhanced cell persistence after irradiation^20^, which may reflect the higher rate of irradiation used in the current study (208 cGy per minute, versus 83 cGy per minute used in prior work), a factor known to alter cell apoptosis in response to irradiation^42^.

**Figure 3:**
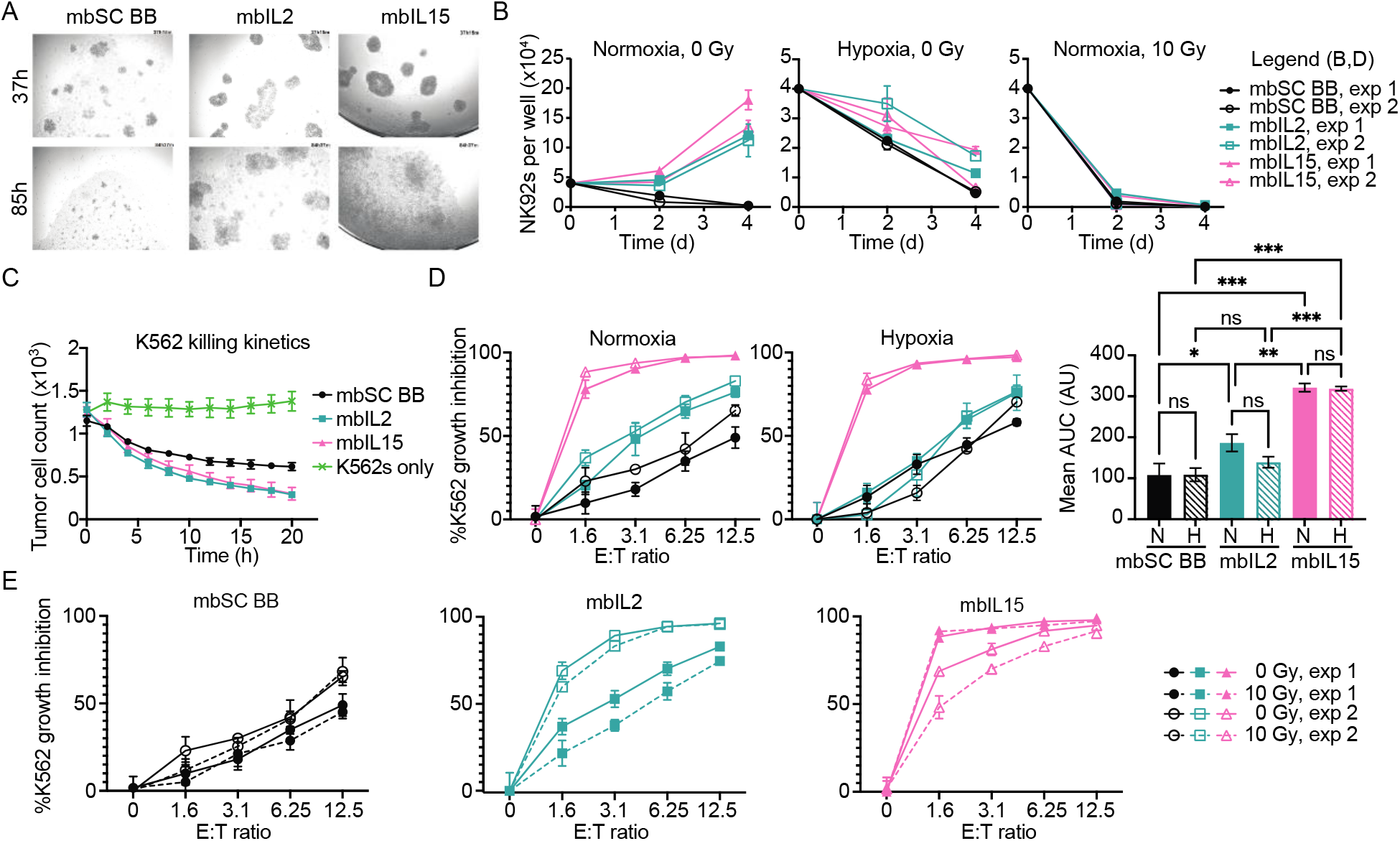
Membrane-bound synthetic cytokine expression enhances the growth and cytotoxicity of NK92 under challenges. **A.** Illustrative images of NK92 growth effects. NK92 transduced to express mbSC were cultured in cytokine-free media and imaged via time lapse microscopy (see also **Supplementary Videos 1-3)**. **B**. Effects of mbSC expression on NK92 growth in normoxia (∼21% O_2_), hypoxia (1% O_2_), or after irradiation (10 Gy). Sample labels in this panel also apply to (**D**). **C.** Effects of mbSC expression on NK92 cytotoxicity against K562. NK92 were cultured without IL2 for 24 h, then mixed with fluorescent K562 at a ratio of 3.125 NK92:1 K562 for 20 h. Tumor cells (mScarlet-I^pos^) were counted using image-based cell-cytometry. **D**. Effects of mbSC expression on NK92 cytotoxicity as a function of effector to target (E:T) ratio and oxygen tension. NK92 were cultured without IL2 for 24 h, then mixed with fluorescent K562 at varying effector (NK92) to target (K562) ratios (E:T ratios) and incubated at oxygen tensions indicated. After 20 h, live K562 counts were obtained by flow cytometry. AUC analysis of K562 inhibition curves was performed. * indicates p<0.05, ** indicates p,0.01, *** indicates p<0.001 (one-way ANOVA,Tukey’s HSD). See **Supplementary Note 1** for full ANOVA results. **E.** Effects of mbSC expression on NK92 cytotoxicity against K562 following irradiation. NK92 were cultured without IL2 for 24 h and irradiated as indicated, then mixed at varying E:T ratios with K562 and incubated in ambient oxygen for 20 h. Live K562 counts were measured by flow cytometry. For all line graphs, points represent means and error bars represent standard deviation, n=3. For all flow cytometry-based experiments, a viability stain was used to exclude dead cells. Abbreviations: arbitrary units (AU), area under the curve (AUC), independent experimental replicate number (exp), hypoxia (H), normoxia (N).

We next examined whether mbSC expression altered NK92 cytotoxicity. For both mbIL2- and mbIL15-expressing NK92 cells, we directly observed cytolysis of K562 B cell lymphoma cells (**Supplementary Videos 4, 6-7**). These cell lines exhibited similar kinetics of activity against K562 over 20 h compared to each other, with both more effective than control NK92 (**Fig. 3C**). Across multiple effector to target (E:T) ratios, when compared to control NK92, both mbSCs significantly enhanced NK92-mediated K562 growth inhibition in normoxia, while in hypoxia, only mbIL15 significantly enhanced K562 growth inhibition (**Fig. 3D**). A possible contributing factor is hypoxia-induced acidification of the media, which reduces IL2 signaling.^43^ Expression of either mbSC evaluated here enhanced NK92 cytotoxicity after cells were irradiated, improving potential clinical utility (**Fig. 3E**). Notably, mbIL2 NK92 showed greater variations in potency against K562 across repeated experiments, which may indicate variations in cell state that are not attributable to known biology and warrant mechanistic investigations in future studies. Altogether, these results show that mbSC expression is a viable strategy for enhancing NK92 growth and cytotoxicity in normoxia and is compatible with irradiation-based protocols to halt NK92 expansion while retaining cytotoxicity.

### Effects of soluble synthetic cytokines on NK92 challenged with hypoxia and irradiation

We next investigated how soluble synthetic cytokine signaling influenced NK92 expansion and cytotoxicity against K562 in various conditions of interest, focusing our investigation on the four soluble synthetic cytokines (sSCs) whose bioactivity was validated in **Fig. 1** (Neo2/15, eIL12, eIL15 and DR18). As recombinant protein monotherapies, these sSCs (or similar constructs) can stimulate NK92 cytotoxicity in vitro (eIL15^30^) and/or activate anti-tumor endogenous immune cell responses in vivo (Neo2/15^24^, eIL12^44^, eIL15^45^, DR18^25^). When produced by NK92, select sSCs can activate proliferation and/or cytotoxicity of co-cultured non-NK92 cells (eIL12^19^, Neo2/15^21^). However, the effects of engineering NK92 to express sSCs on their own proliferation and cytotoxicity are not well defined. To address this gap in knowledge, we systematically tested how sSC expression affected NK92 expansion and cytotoxicity, including evaluations in hypoxia and after irradiation. Based on the effects of their natural analogs and prior literature, we expected that, in standard conditions, Neo2/15 and eIL15 would enhance growth and cytotoxicity^21,33^, while eIL12 and DR18 would only enhance cytotoxicity^11,34,37^. We further expected hypoxia and irradiation to limit the effects of sSC expression on growth and cytotoxicity.^46,47^

We first tested whether sSC expression induced NK92 expansion. When cultured without IL2 in normoxia, Neo2/15 expression led to the highest cell counts after incubation in normoxia; eIL15 also supported NK92 expansion to a lesser degree, which might reflect different affinities of the two cytokines for their receptor (**Fig. 4**). No sSC improved NK92 persistence or growth in hypoxia by day 4, though some permitted mildly improved survival on day 2 (**Fig. 4**). All sSC-expressing cell lines remained sensitive to irradiation (**Fig. 4**), although sp^IgE^Neo2/15 conferred marginally higher survival (**Fig. 4A**).

**Figure 4:**
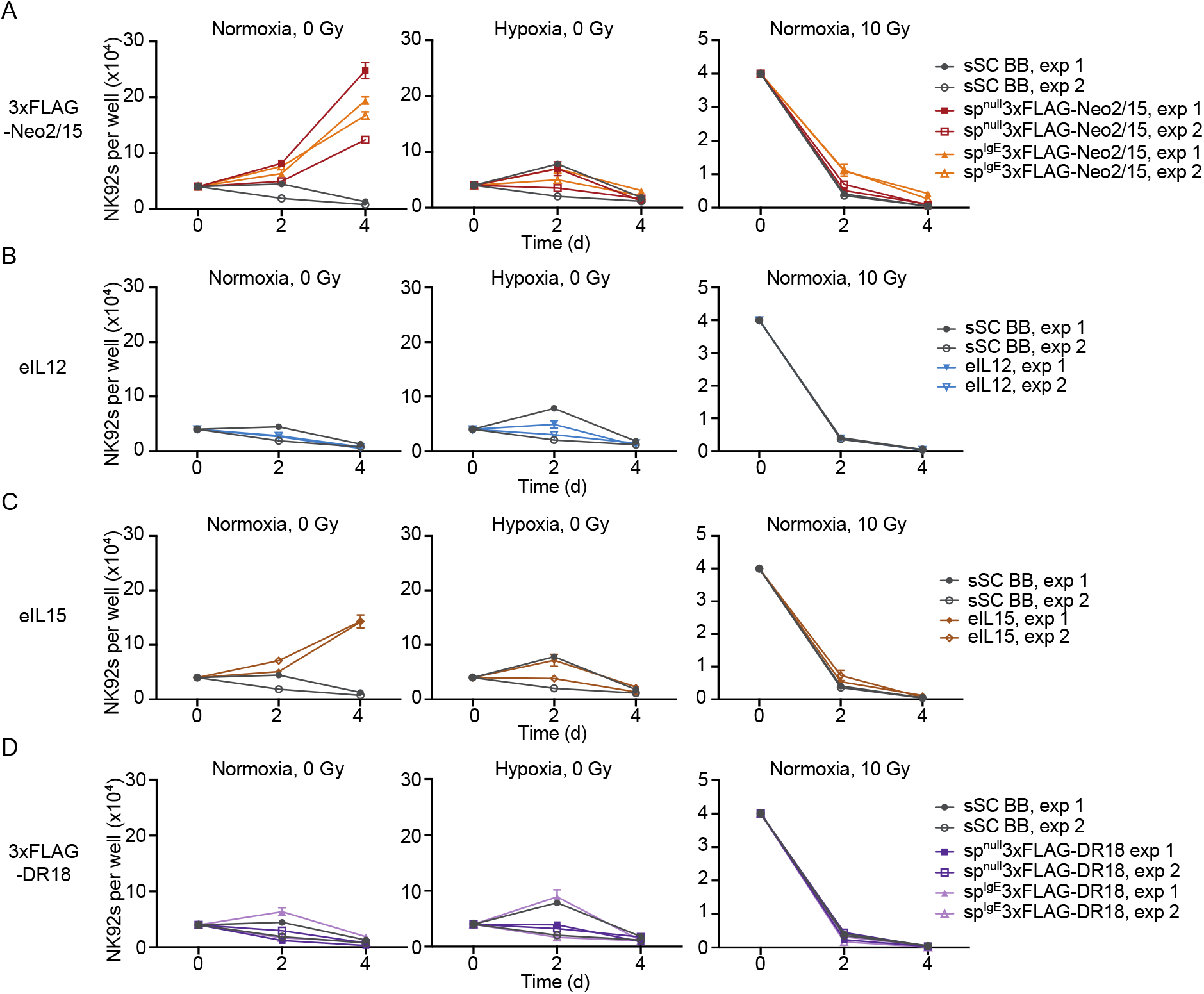
Soluble synthetic cytokine expression modulates the growth of NK92 under challenges. **A-D.** Effects of sSC expression on NK92 growth in ambient oxygen (∼21% O_2_) or hypoxia (1% O_2_) or after irradiation (10 Gy) (and then grown in ambient oxygen). For all line graphs, points represent means and error bars represent standard deviation, n=3. To aid in comparison, growth of sSC BB NK92 was plotted on A-D, but represent the same set of experiments. Abbreviations: cells transduced with an empty sSC backbone expression vector (sSC BB); cells transduced with the indicated sSC lacking a signal peptide (sp^null^); cells transduced with the indicated sSC including an IgE signal peptide (sp^IgE^).

We next investigated how sSC expression modulates NK92 cytotoxicity. NK92-mediated cytolysis was visualized using time-lapse microscopy for each sSC-expressing NK92 line to validate that engineered NK92 remained cytotoxic towards K562 (**Supplementary Videos 4, 8-14**). Across multiple effector to target (E:T) ratios, sSC-expression increased NK92-mediated K562 growth inhibition in both normoxia and hypoxia (**Fig. 5A-E**). There was some variation in rate of K562 killing in normoxia, with eIL12, sp^null^ and sp^IgE^Neo2/15 expression leading to the most rapid and prolonged cytotoxicity (**Fig. 5F**). Irradiation with 10 Gy did not diminish the effects of sSC expression on NK92-mediated K562 growth inhibition (**Fig. 5G**). Altogether, these observations indicate that, as expected, sSCs analogous to IL2 or IL15 enhance NK92 growth, unless cells are challenged with irradiation or hypoxia, and that, excitingly, all sSCs enhance NK92 cytotoxicity, even when cells are challenged with irradiation or hypoxia.

**Figure 5:**
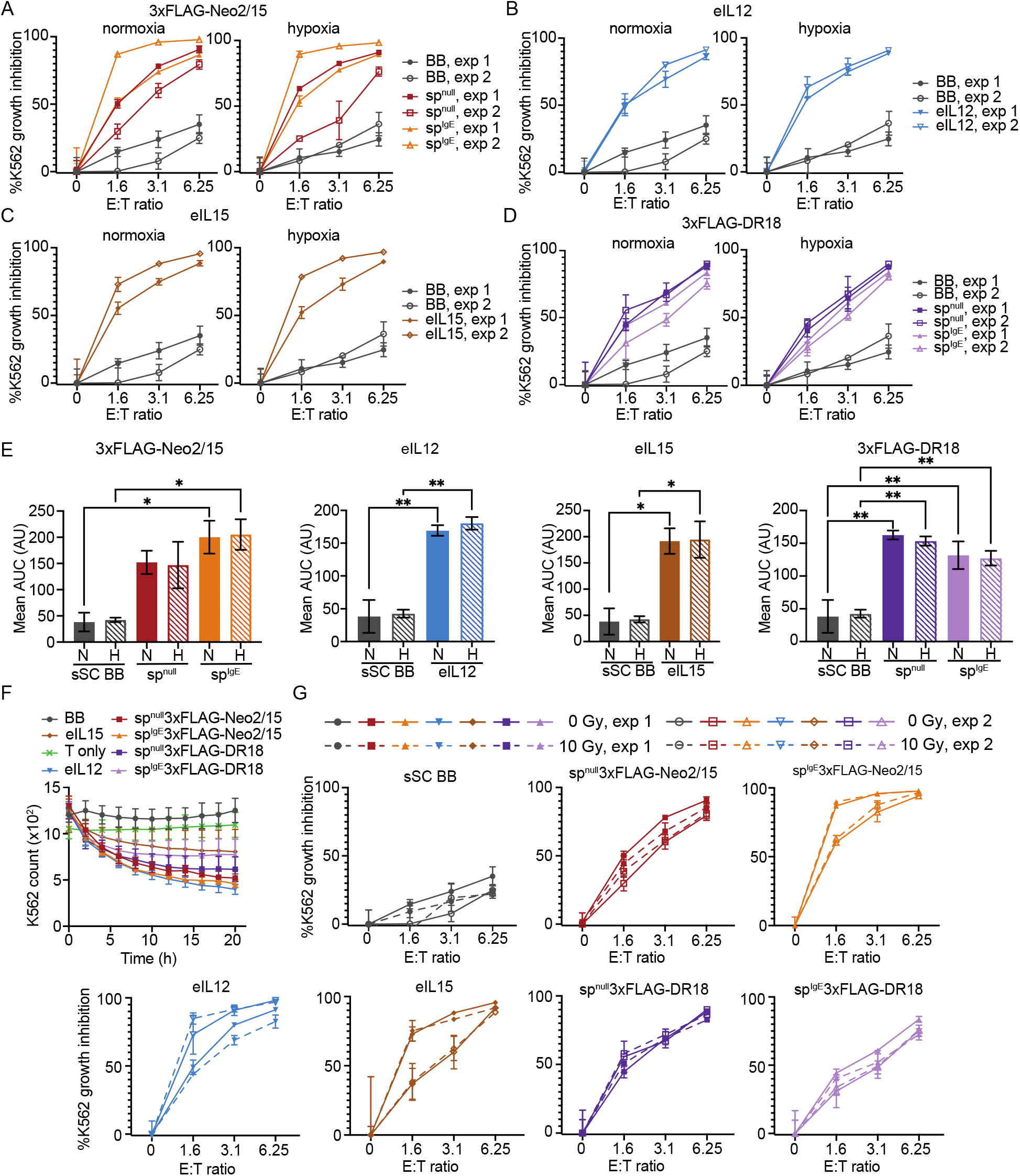
Soluble synthetic cytokine expression modulates the cytotoxicity of NK92 under challenges. **A-E**. Effects of sSC expression on NK92 cytoxocity as a function of effector to target (E:T) ratio and oxygen tension. NK92 were cultured without IL2 for 24 h, then mixed with fluorescent K562 at varying effector (NK92) to target (K562) ratios (E:T ratios) and incubated at oxygen tensions indicated. After 20 h, live K562 counts were obtained by flow cytometry. Panel (**E**) shows analyses of corresponding E:T data series. * indicates p<0.05, ** indicates p<0.01 (one-way ANOVA,Tukey’s HSD). See **Supplementary Note 1** for full ANOVA results. **F.** Effects of sSC expression on rate of NK92 cytotoxicity against K562. NK92 were cultured without IL2 for 24 h, then mixed with fluorescent K562 at a ratio of 3.125 NK92:1 K562 for 20 h. Tumor cells (mScarlet-I^pos^) were counted using image-based cell-cytometry. **G**. Effects of sSC expression on NK92 cytotoxicity against K562 following irradiation. NK92 were cultured without IL2 for 24 h and irradiated as indicated, then mixed at varying E:T ratios with K562 and incubated in ambient oxygen for 20 h. Live K562 counts were measured by flow cytometry. For all line graphs, points represent means and error bars represent standard deviation, n=3. For all flow-cytometry based experiments, a viability stain was used to exclude dead cells. Abbreviations: arbitrary units (AU), area under the curve (AUC), independent experimental replicate number (exp), cells transduced with an empty backbone expression vector (BB); cells transduced with the indicated sSC lacking a signal peptide (sp^null^); cells transduced with the indicated sSC including an IgE signal peptide (sp^IgE^).

### Paracrine effects of synthetic cytokine expression on NK92 growth and cytotoxicity

We next investigated whether synthetic cytokine (SC) expression confers paracrine signaling that is sufficient to modulate NK92 growth or cytotoxicity. Paracrine signaling plays an important role in the efficacy of cell-based therapies.^48^ Paracrine activity can be cell contact-dependent (e.g., cytokine-receptor trans-presentation)^49,50^ or contact-independent (e.g., secretion and diffusion of soluble cytokines). Based on the literature and data in Figures 4 and 5, we expected soluble SCs to confer appreciable contact-independent paracrine effects on NK92 growth and/or cytotoxicity that for Neo2/15 and DR18 would be enhanced by the inclusion of IgE signal peptides. As mbIL2 and mbIL15 are membrane tethered and non-transduced cells survived despite cytokine withdrawal when co-cultured with mbIL2 NK92, but not mbIL15 NK92 (**Fig. 2D**), we also expected mbIL2, but not mbIL15, to have contact-dependent paracrine effects.

We first investigated paracrine effects upon NK92 growth. Because expression of mbIL2, mbIL15, Neo2/15 (with and without IgE signal peptide), and eIL15 promoted growth in NK92 producer cells, the paracrine activity of these factors was characterized by evaluating the ability to promote growth of co-cultured NK92 expressing no synthetic cytokines (SC^neg^ NK92). All SC-expressing NK92, including mbIL15 NK92, conferred appreciable paracrine effects on SC^neg^ NK92 growth (**Fig. 6A**). Given this surprising observation with mbIL15, we speculate that during cytokine-withdrawal selection of mbIL15 NK92 (**Fig. 2C**), it is possible that non-transduced cells were not detected because the autocrine mbIL15 signaling potency is stronger than its paracrine activity and this enabled transduced cells to outcompete non-transduced cells, or that the magnitude of paracrine signaling was lower in these prior experiments. To test whether the observed paracrine signaling is contact-dependent, cell media conditioned by SC-expressing NK92 was tested for the ability to induce SC^neg^ NK92 expansion. Interestingly, despite being membrane-tethered, expression of both mbIL2 and mbIL15 conferred contact-independent effects (**Fig. 6B**). We speculated that this effect could be mediated by cleavage-mediated release of mbSC components, by mbSC-induced secretion of other factors that can support NK92 growth in trans, and/or by extracellular vesicle (EV)-mediated transfer of mbSCs, as EVs are able to mediate signaling between immune cells^51^ (these possibilities are further investigated in subsequent experiments). Conditioned medium from sp^IgE^Neo2/15 and eIL15 NK92, but not sp^null^Neo2/15, also significantly supported SC^neg^ NK92 expansion (**Fig. 6B**). Sp^null^Neo2/15 might be secreted via an unconventional pathway^52^ at a level insufficient to support SC^neg^ cell growth in the absence of cytokine-producing cells, although it remains possible that the paracrine effects of sp^null^Neo2/15 are contact-dependent.

**Figure 6:**
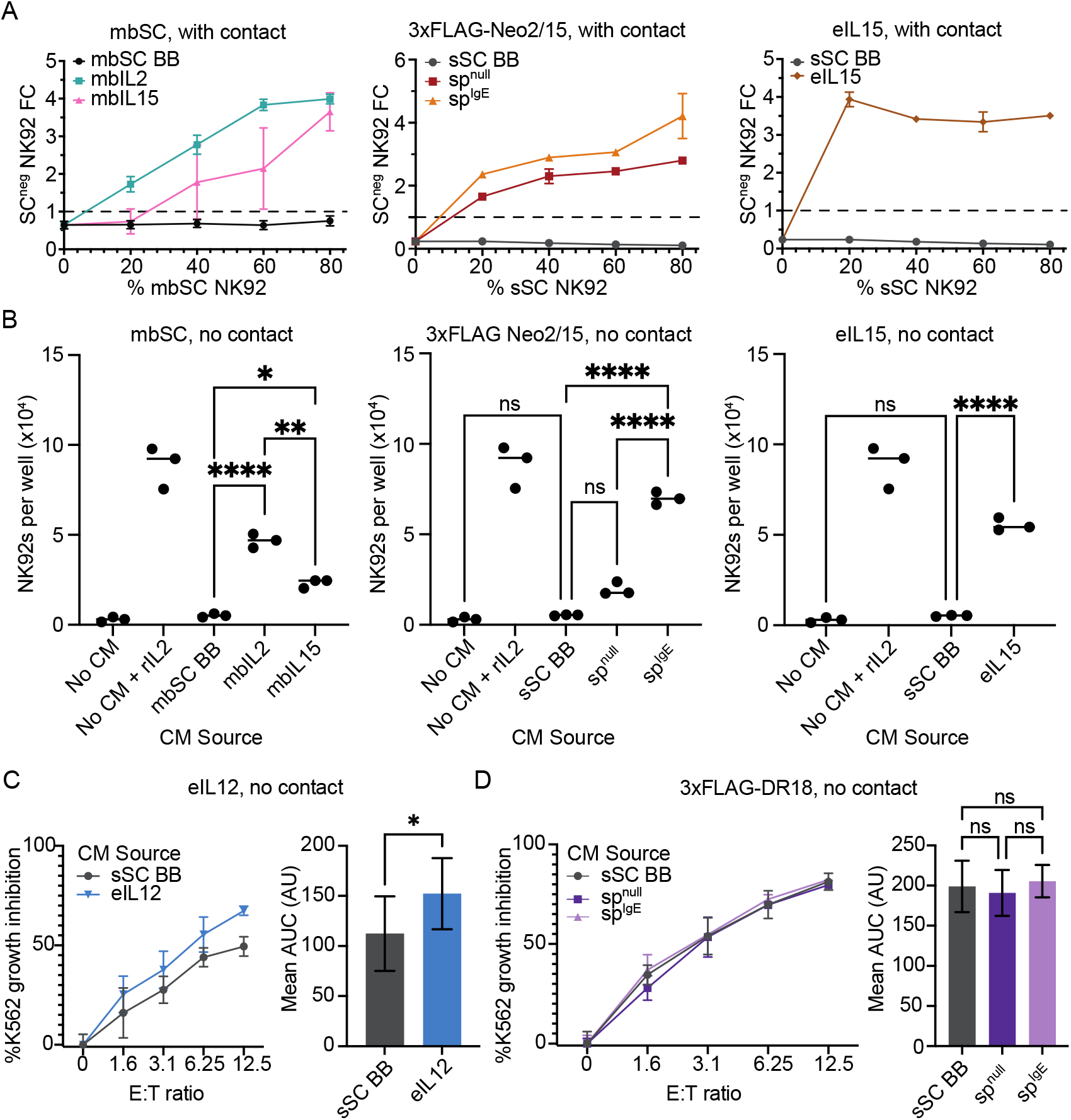
Select synthetic cytokines are sufficient to exert paracrine effects on NK92 growth or cytotoxicity. **A.** Paracrine effects of SCs on SC^neg^ NK92 growth with cell contact. SC NK92 were co-cultured for 3 d with SC^neg^ NK92, after which SC^neg^ NK92 were counted by flow cytometry. SC^neg^ NK92 fold change was calculated by dividing SC^neg^ NK92 count by number of SC^neg^ NK92 seeded. **B.** Paracrine effects of SCs on SC^neg^ NK92 growth without cell contact. Media were conditioned for 4 d by SC NK92, then clarified to remove cells. SC^neg^ NK92 were cultured in 50% conditioned media for 3 d, then counted by flow cytometry. rIL2 indicates 100 IU/mL recombinant IL2. Control cell data points are duplicated across panel. **C-D.** Paracrine effects of SCs on SC^neg^ NK92 cytotoxicity without cell contact. Media was conditioned by eIL12 (**C**) or 3xFLAG-DR18 (**D**) NK92 for 1 d, then clarified to remove cells. As 5 IU/mL recombinant IL2 enhanced effects of DR18 on NK92 cytotoxicity (Fig. 1K), 5 IU/mL recombinant IL2 was added to all conditioned media conditions in (**D**), only. SC^neg^ NK92 were incubated with fluorescent K562 (mScarlet-I^pos^) for 20 h in 50% conditioned medium, after which tumor cells were counted by flow cytometry. K562 growth inhibition curves were generated and analyzed. In line graphs, points represent mean and error bars represent standard deviation. n=3. Bar height indicates mean AUC, error bars indicate SEM. For all experiments, a cell viability stain was used to exclude dead cells from analysis. ns indicates p>0.05, * indicates p<0.05, ** indicates p<0.01, and **** indicates p<0.0001 as calculated by a two-tailed t-test (panel (**C**) only), or an one-way ANOVA followed by Tukey’s HSD (all other panels). See **Supplementary Note 1** for full ANOVA results. Abbreviations: arbitrary units (AU), area under the curve (AUC), fold change (FC), conditioned medium (CM), standard error of the mean (SEM), cells transduced with an empty backbone expression vector for either membrane bound synthetic cytokines (mbSC BB) or soluble synthetic cytokines (sSC BB); cells transduced with the indicated sSC lacking a signal peptide (sp^null^); cells transduced with the indicated sSC including an IgE signal peptide (sp^IgE^).

We next investigated paracrine effects upon NK92 cytotoxicity. Because expression of DR18 and eIL12 signaling enhances cytotoxicity of NK92 expressing these cytokines, we investigated paracrine activity of these factors vis-à-vis enhancement of SC^neg^ cytotoxicity against K562. Since we could not deconvolute K562 killing by SC and SC^neg^ NK92 in a mixed co-culture format, which would be necessary for assessing contact-dependent paracrine effects, only contact-independent paracrine effects were evaluated. Conditioned medium from eIL12 NK92, but not DR18 (regardless of signal peptide), significantly enhanced the SC^neg^ NK92-mediated K562 growth inhibition (**Fig. 6C-D**). It is possible that DR18 signaling is sufficient for priming NK92 cytotoxicity over long exposures but is insufficient for rapid paracrine activation of NK92 cytotoxicity in the absence of other cytokines. DR18 might also act by prolonging the pro-cytotoxic effects of recombinant IL2 after it is withdrawn. Taken together, these observations indicate that paracrine effects of SC expression in NK92 are substantial, varied, and sometimes surprising given prior knowledge, motivating subsequent mechanistic investigations.

### Mechanistic evaluation of paracrine effects mediated by membrane-bound synthetic cytokines

To elucidate how the expression of membrane-bound synthetic cytokines (mbSCs) conferred contact-independent paracrine effects, we next investigated whether these effects could be mediated by extracellular vesicles (EVs), non-EV soluble factors, or both (**Fig. 7**). EVs are a heterogeneous group of membrane-bound vesicles produced by virtually all cell types.^53^ EVs present some membrane proteins on their surface and can encapsulate cytosolic proteins; the composition of EVs remains an active area of investigation.^54^ The importance of EVs in intercellular communication, including communication between immune cells, is increasingly recognized.^51^ To explain the surprising aforementioned observations (**Fig. 6B**), we hypothesized that mbSC signaling could induce NK92 to secrete soluble factors that support growth in trans, mbSCs could be cleaved at the membrane to release soluble cytokines, and/or mbSCs could be packaged onto EVs and secreted to subsequently stimulate recipient NK92 (**Fig. 7A**).

**Figure 7:**
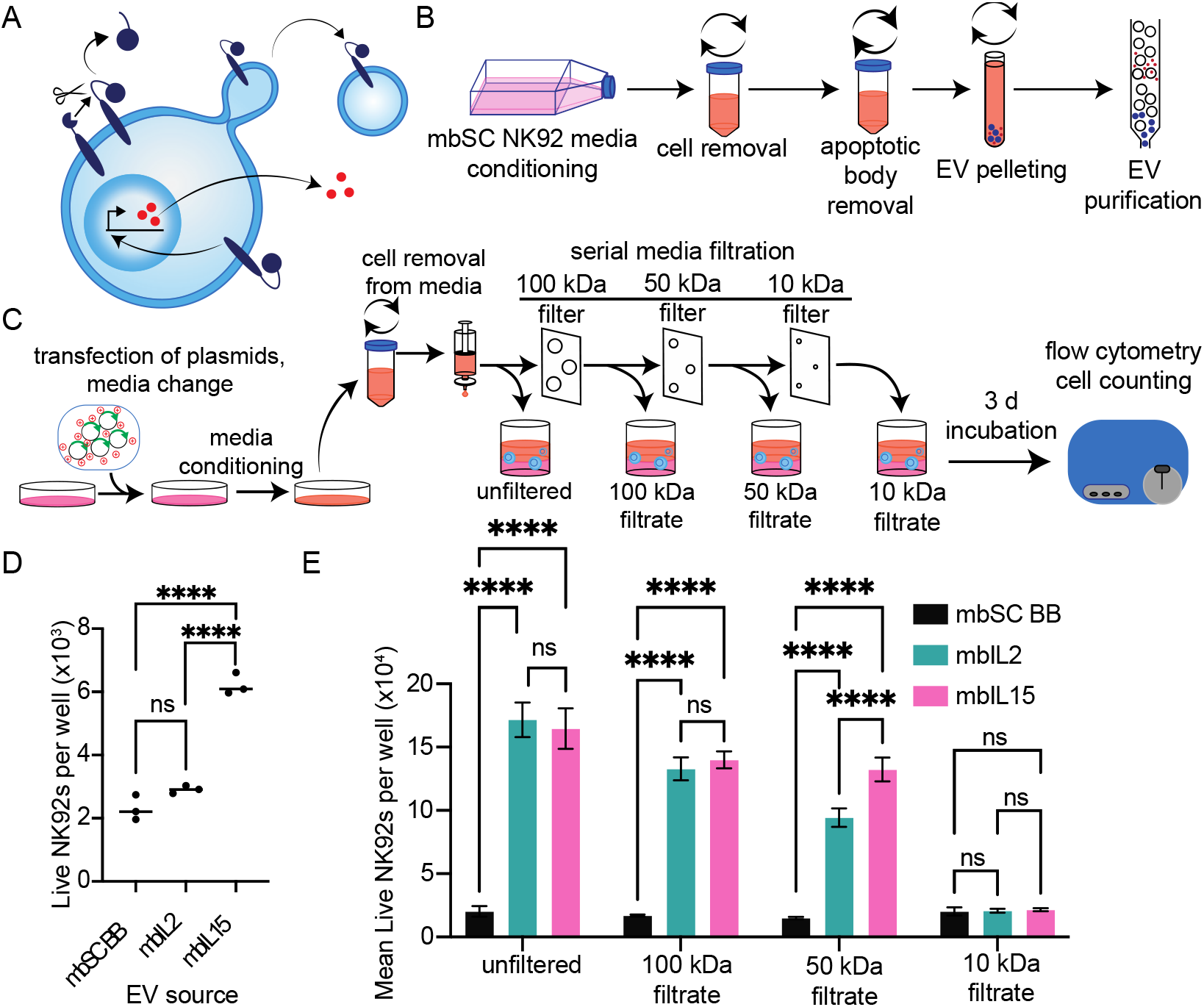
EVs partly mediate mbIL15 paracrine activity and non-EV components mediate paracrine activity of both mbIL2 and mbIL15. **A.** Schematic of possible mechanisms by which mbSC expression mediates paracrine effects on NK92 growth. **B.** Schematic of EV isolation from mbSC NK92 **C.** Schematic of assay used to test how size-based depletion of soluble factors from medium conditioned by mbSC-expressing HEK293FTs affects the medium’s paracrine effects on NK92 growth. **D.** EV-mediated paracrine support of NK92 growth. Serum-free medium was conditioned by mbSC-expressing NK92 for 2 d. EVs were isolated from conditioned media and added to SC^neg^ NK92 (3.7×10^9^ EVs per well). After 3 d, live SC^neg^ NK92 were counted by flow cytometry. **E.** Evaluation of sub-EV contributors to paracrine effects attributable to mbSC expression. HEK293FTs were transfected with the mbSCs, and media were conditioned for 24 h before depleting EVs and other species using serial filtration through 100 kDa, 50 kDa, and finally 10 kDa molecular weight cutoff filters. The flow through from each filtration step was evaluated for the ability to support SC^neg^ NK92 growth; SC^neg^ NK92 were cultured in 50% filtered conditioned media for 3 d, then quantified by flow cytometry. Error bars indicate standard deviation. For all experiments, **** indicates p<0.0001 (one-way (**D**) or two-way (**E**) ANOVA followed by Tukey’s HSD), n=3. See **Supplementary Note 1** for full ANOVA results. For all experiments, a viability stain was used to exclude dead cells. Abbreviations: extracellular vesicle (EV), cells transduced with an empty membrane bound cytokine backbone expression vector (mbSC BB), kilodalton (kDa).

We first evaluated whether EVs from mbSC-expressing NK92 might contribute to the observed paracrine effects. EVs were isolated from serum-free medium conditioned by mbSC NK92 using ultracentrifugation, followed by size-exclusion chromatography (**Fig. 7B**). Isolated EVs exhibited appropriate sizes, protein markers, and morphology expected for EVs according to best practice guidelines as published in the EV literature^55^, validating the EV isolation methodology (**Supplementary Fig. 2B-D**). EVs from mbIL15 NK92, but not mbIL2 NK92, were sufficient to support SC^neg^ NK92 growth (**Fig. 7D**). This could reflect differential trafficking of the mbSC transmembrane domains (IL15Rα versus IL2Rβ, respectively) to NK92 EVs, or differential potency of these factors in this format.

We next investigated whether soluble factors smaller than EVs mediated mbSC paracrine effects. To test this possibility, conditioned media was passed through filters with progressively smaller molecular weight cutoffs (MWCO) and evaluated for the potential to support SC^neg^ NK92 growth. We decided to focus specifically on evaluating the potential of mbSC expression to confer effects mediated by mbSC components (rather than mbSC-expression-induced secretion of other factors by NK92), and for that reason we evaluated media conditioned by transiently transfected HEK293FTs (which should not express receptors for these factors). To deplete EVs, conditioned media were ultrafiltered through a 100 kDa MWCO filter, which is sufficient to remove EVs^56^. To assess whether a lower limit in the size of bioactive compounds could be identified, media were then passed through a 50 kDa MWCO filter and a 10 kDa MWCO filter, which would remove free IL2 (∼16 kDa) and IL15 (∼15 kDa) (**Fig. 7C**). For both mbSCs, non-EV factors were sufficient to induce SC^neg^ NK92 growth, and these factors were greater than 10 kDa in size (**Fig. 7E**). These factors may be mbSC cleavage products, as the transmembrane domains of both mbIL15 and mbIL2 are known to be cleaved at the membrane.^57,58^ Intriguingly, mbIL2-conditioned media was significantly less effective than was mbIL15 conditioned media after filtration through a 50 kDa MWCO filter (**Fig. 7E**). This may reflect the removal of bioactive soluble mbIL2 multimers by that filter.^59^ In sum, these observations indicate that paracrine activity of mbSC expression can be mediated by EVs as well as secreted subunits of these transgene products, and these effects vary by mbSC identity.

### Evaluating the potential of combined synthetic cytokine expression for engineering NK92

NK92 cell-based therapies that combine the expression a membrane-bound synthetic cytokine (mbSC) and soluble synthetic cytokine (sSC) may benefit from both potent mbSC autocrine activation and delivery of sSCs to modulate endogenous immune cells. However, sustained cytokine overexposure can also lead to NK cell dysfunction^60^. Therefore, these tradeoffs must be evaluated to identify suitable combinations and even interaction effects. As a first step towards evaluating how exposure to signaling from both an mbSC and sSC impacts NK92 expansion and cytotoxicity, we employed a consortia model in which populations of mbSC-expressing NK92 and sSC-expressing NK92 were mixed and evaluated.

We tested how mixed consortia of SC-expressing NK92 modulate expansion and cytotoxicity. Consortia of NK92 were incubated for 3 d, after which each subpopulation’s cell count was measured (**Fig. 8A-B**). NK92 expressing mbIL2 or mbIL15 did not exhibit altered growth when mixed into consortia with NK92 expressing Neo2/15 or eIL15 (**Fig. 8A**), and the converse was also true (**Fig. 8B**). NK92 expressing eIL12 or DR18 exhibited greater expansion when co-cultured with mbIL2- or mbIL15-expressing NK92 (**Fig. 8B**). Similarly, mbIL15-expressing NK92 exhibited slightly (but significantly) increased expansion when cultured with sp^IgE^3xFLAG-DR18 NK92 (**Fig. 8A**), which may reflect synergistic effects of the two cytokines on proliferation^61^. Overall, NK92 expansion was not diminished by (and in some cases was increased by) exposure to multiple SCs. We observed a similar pattern in evaluations of cytotoxicity, in that mixing mbSC NK92 and sSC NK92 into consortia did not diminish the overall population cytotoxicity (**Fig. 8C**). Notably, SC lines showed varying capacity to rescue population cytotoxicity when in consortia with SC^neg^ NK92 cells. In particular, mbIL2 and eIL15 NK92 fully rescued population cytotoxicity, as their cytotoxicity as pure populations did not differ significantly from their cytotoxicity when in consortia with SC^neg^ NK92. Altogether, these consortia analyses suggested that combinatorial engineering of NK92 could be feasible.

**Figure 8:**
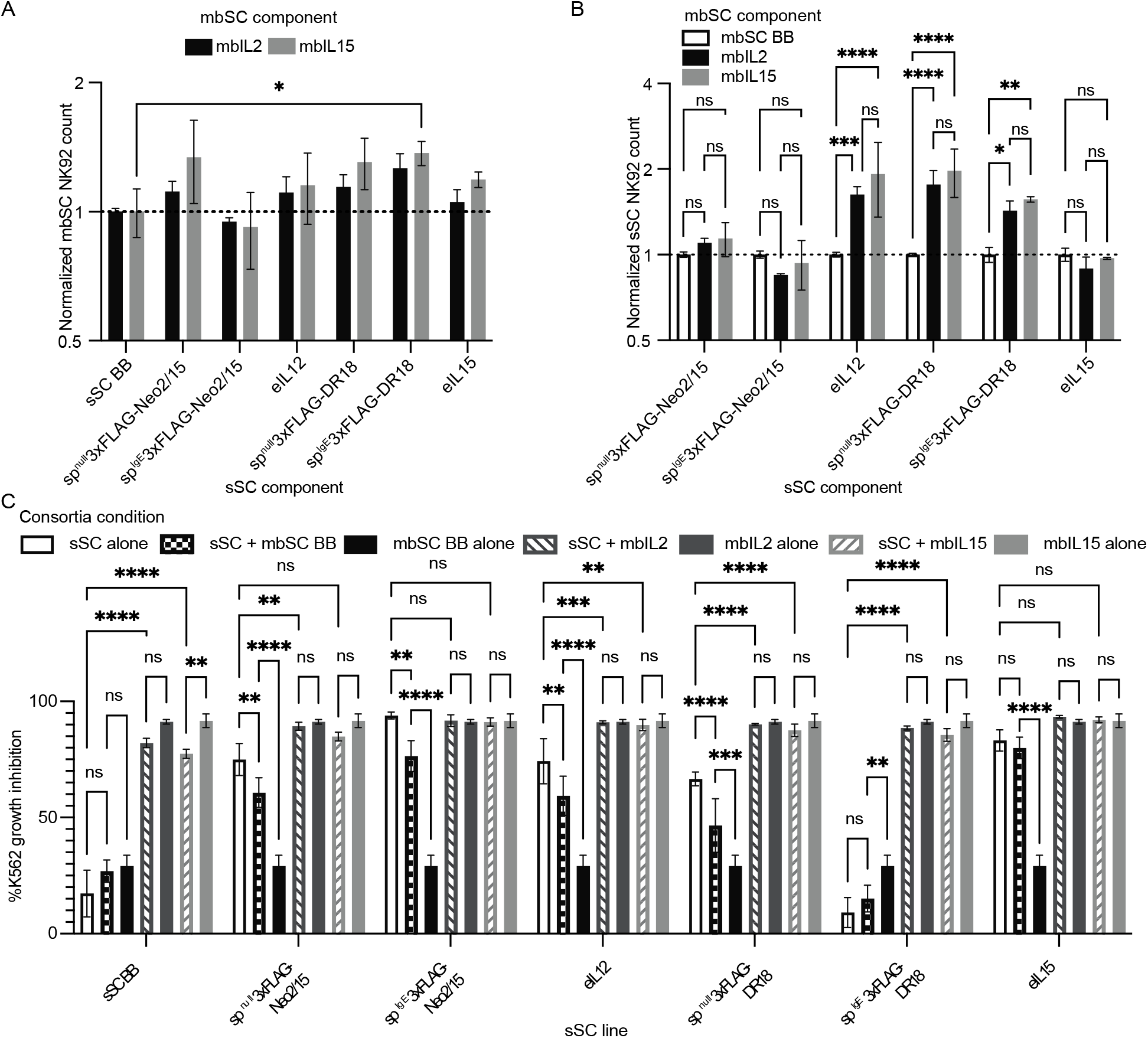
Synthetic cytokine-engineered NK92 consortia enhance growth and cytotoxicity: **A-B.** Expansion of SC-expressing NK92 when in consortia. MbSC NK92 were co-cultured with sSC NK92 at a 1:1 ratio for 3 d, after which mbSC NK92 (miRFP720^pos^) (**A**) and sSC NK92 (mNeonGreen^pos^) (**B**) were counted by flow cytometry. Normalized count was calculated by dividing SC NK92 counts by the average SC NK92 count when co-cultured with BB NK92; see **Equation 2** (Methods). Bar height indicates mean normalized count. Error bars represent standard deviation. n=3. **C.** sSC NK92 alone, or mixed 1:1 with mbSC NK92, were co-incubated with fluorescent K562 cells at a ratio of 3.125:1 NK92:K562 for 20 h. Live K562 (mScarlet-I^pos^) were counted by flow cytometry and used to calculate %K562 growth inhibition. Bar height indicates mean %K562 growth inhibition. Error bars indicate standard deviation. n=3. For all experiments, a viability stain was used to exclude dead cells. ns indicates p > 0.05, * indicates p < 0.05, ** indicates p < 0.01, *** indicates p < 0.001, and **** indicates p < 0.0001 (two-way ANOVA followed by Tukey’s HSD). n=3. See **Supplementary Note 1** for full ANOVA results. Abbreviations: cells transduced with an empty backbone expression vector for sSCs (sSC BB) or mbSCs (mbSC BB).

## Discussion

The findings presented in this study have several implications for engineering NK92 to express synthetic cytokines and will inform future efforts to design NK92 cell therapies, which may expand the availability of cell-based cancer therapies to more patients. Although prior investigations have evaluated expression of some synthetic cytokines in NK92 to enhance their function, no prior analysis includes comparative evaluation across properties important for cell therapy production and performance. After platform development focused on enabling or enhancing expression of SCs, our evaluation included properties useful for both cell therapy production (selection of engineered cells, expansion in cytokine-free media) and performance (cytotoxicity, including after exposure to irradiation and hypoxia). Finally, we generated a series of new insights into the mechanisms by which SC-expressing NK92 may function in paracrine fashion, and in which combinatorial SC effects may drive NK92 growth and performance.

This study generated multiple novel insights important to enabling or enhancing expression of SCs. We established, for the first time, that DR18 and single-chain IL12, the latter only when expressed at attenuated levels, can be expressed constitutively by NK92 to a degree sufficient to increase NK92 cytotoxicity. The feasibility of constitutively expressing DR18 and eIL12 was surprising, as prolonged exposure to their native analogs induce NK92 death^38,39^. It is possible that low level IL12 expression leads to levels of IL12 signaling closer to those experienced in vivo or that IL12 exerts distinct NK92 responses as a function of dose. We also found that Neo2/15 is highly sensitive to C-terminal modification and that and shows appreciable autocrine activity despite lacking a known secretion tag, though addition of an IgE signal peptide substantially enhanced its effects. Similarly, DR18 showed appreciable autocrine activity despite lacking a secretion tag. This is surprising for Neo2/15, as it is a fully synthetic amino acid sequence that lacks known signals for secretion, while this is less surprising for DR18, as its sequence is derived from IL18, which is secreted via an unconventional pathway dependent on plasma membrane permeability^62^.

This study also showed, for first time, that SCs impart substantially different selective pressures on NK92. It was unsurprising that mbSCs, but not SCs, positively selected for NK92 when they were cultured in cytokine-free media, as mbIL2 and mbIL15 were known to preferentially support transduced NK92 growth ^20,22^, while soluble SCs were expected to support the growth of both transduced and non-transduced cells. However, it was surprising that mbIL2 selected for lower expression loci than mbIL15. One possibility is that mbIL2 activates NK92 growth signaling more potently than does mbIL15, enabling survival of NK92 with lower transgene expression. Another possibility is that mbIL2 is toxic above certain expression levels, negatively selecting cells with high transgene expression loci. Regardless, the different selection mediated by these mbSCs is particularly surprising as the IL2 and IL15 receptors only differ in their alpha subunits, which are not thought to contribute to cytokine signaling, though the contribution of IL15Rα to IL15 signaling is becoming more appreciated.^63^ It is possible that signaling from IL15Rα in mbIL15 is responsible for the different mbSC selection patterns. It is also possible that mbIL2 and mbIL15 complex with other signaling chains with different affinity, leading to differential signaling. Finally, it is possible that mbIL2 and mbIL15 have different stabilities, necessitating different rates of expression for the same level of surface expression. It was also surprising that Neo2/15 exerted negative selective pressures, which is likely due to exhaustion from high cytokine signaling^64^ or cell burden due to transgenic protein production.

This study also generated insights into the impacts of SC expression on NK92 performance. As expected, SCs analogous to IL2 or IL15 supported NK92 expansion, but not when challenged with hypoxia or irradiation. Excitingly, all SCs but mbIL2 retained effects on cytotoxicity in hypoxia, and all SCs retained their effects on cytotoxicity when irradiated. The discrepancy between growth and cytotoxicity might be due to disruption of distinct signaling mechanisms that mediate growth but not cytotoxicity, or may reflect that SC enhancement of cytotoxicity, but not expansion, is the result of being primed with cytokine signaling, rather than cytokine signaling concurrent with cytolysis. Regardless, the SCs increase the cytotoxicity of NK92 without mitigating their growth inhibition by irradiation, which mitigates concern that these SCs may override this critical safety process. The loss of mbIL2 effects on cytotoxicity in hypoxia was unexpected and might reflect induction of a state that is more sensitive to hypoxia.

This study additionally revealed insights into the paracrine effects of mbIL2 and mbIL15. Both mbSCs had contact-independent paracrine effects on NK92 expansion. These findings are likely due, in part, to receptor chain cleavage and release of soluble cytokine-cytokine receptor complexes, which have been described for both IL2-IL2Rβ^59^ and IL15-IL15Rα^58^ complexes, the cytokine-cytokine receptor pairs used in mbIL2 and mbIL15, respectively. Notably, these prior investigations, as well as our study of sub-EV mediators of mbSC paracrine effects, did not express cytokines in NK92, and it remains possible that these findings do not translate to paracrine effects of mbSCs produced by NK92 (e.g., due to differential expression of proteases responsible for cytokine receptor cleavage). We also discovered that extracellular vesicles were a novel mechanism by which mbIL15 exerts paracrine effects, and that, surprisingly, this mechanism did not also apply to mbIL2. Extracellular vesicles isolated from mbIL15, but not mbIL2, expressing NK92 were sufficient to induce NK92 growth. There are multiple possible explanations for how mbIL15 mediates effects via vesicles. It might be presented on the surface of vesicles and directly activate IL15 receptors on NK92. It is also possible that vesicles are endocytosed and receptors are trafficked to a suitable membrane location in the recipient cell, where they influence signaling. The unexpected difference in the role of EVs in mediating paracrine effects of mbIL2 versus mbIL15 may be due to differential localization within intracellular membrane compartments^65,66^, which might affect their relative propensity to be packaged into vesicles.

Finally, our consortium analyses indicate that signaling from membrane-bound cytokines and soluble synthetic cytokines can be combined without compromising NK92 expansion or cytotoxicity, and such strategies even enhance the expansion of NK92 expressing DR18 or eIL12. This finding supports the potential utility of developing NK92 therapies that combine such modalities. Combining modalities is an attractive design because mbSCs enable production and performance of engineered NK92 therapies, while sSCs have previously demonstrated abilities to activate antitumor responses by endogenous immune cells or co-administered adoptive cell therapies^19,23-26^. While the findings of our consortium analysis were largely expected, the outcomes were nonetheless non-obvious, as it was possible that cytokine overstimulation would lead to cell dysfunction or death. These analyses also provide a useful reference point for future efforts to engineer NK92 with a single vector that combines expression of mbSCs with sSCs, which may be complicated by competing selective pressures, resource competition from high protein expression in a single cell, or effects of prolonged exposure to more than one cytokine. Overall, the findings described herein contributed multiple new biological insights into the behavior and effects of SCs expressed in NK92, which will enable future engineering efforts that leverage SCs to enhance NK92 therapy production and performance.

## Conclusions

This comparative analysis of the effects and interactions of synthetic cytokine expression may guide NK92 bioengineering and improve the feasibility and function of NK92 cell-based therapies. This study generated a suite of hypotheses that may guide future validation work relating in vitro performance to in vivo performance, which is most appropriately deployed across a range of cancer models, as performance varies by model and disease. Another frontier is the combination of these synthetic cytokine strategies with other functional transgene (e.g., expression of CARs to enhance tumor-reactivity). While this study focused on NK92 therapies, the framework for comparatively evaluating synthetic cytokines may be extended to evaluate other cell-based therapies. Ultimately, this analysis revealed multiple avenues for continuing to improve NK92 bioengineering towards realizing the potential of effective, off-the-shelf anti-cancer cell therapeutics.

## Online Methods

### General DNA assembly and synthetic cytokine modification

Plasmid construction was completed using standard molecular biology techniques. Polymerase chain reactions were completed using Phusion DNA Polymerase (New England Biolabs). Plasmids were assembled using restriction enzyme cloning. Transgenes were codon-optimized for production in human cells, using the GeneArt Optimizer (ThermoFisher). A (GGGGS)_3_ linker was inserted between the two IL12 monomers to enhance protein stability^67^. A 3xFLAG tag followed by a GSG linker was added to the N terminus of Neo2/15. A 3xFLAG tag followed by a GSG linker was added to the N terminus of DR18. The FLAG epitope tag was removed between the signal peptide and coding sequences of eIL15. Plasmids created in this study are deposited with and distributed by Addgene at https://www.addgene.org/Joshua_Leonard/. Plasmid sequences for constructs created in this study are included in supplementary material online.

### Cell culture

HEK293FT (Thermo Fisher R70007) and HEK293T Lenti-X (Takara Bio #632180) were cultured in Dulbecco’s Modified Eagle Medium (Gibco #31600-091) supplemented with 10% heat-inactivated fetal bovine serum (Gibco #16140-071), 100 U/mL penicillin, 100 µg/mL streptomycin (Gibco #15140122), and 4mM L-glutamine (Gibco #25030-081). For Lenti-X cells, medium was supplemented with 1 mM sodium pyruvate (Gibco #11360070). To split HEK293FT and Lenti-X cells, spent medium was aspirated, cells were rinsed with 5 mL of sterile phosphate buffered saline then incubated with 1.5 mL of Trypsin-EDTA at 37°C for 2-5 min. Cells were resuspended in fresh media, then plated in 10 cm dishes. K562 B-cell lymphoma cells (ATCC, CCL-243) were cultured in Iscove’s Modified Eagle Medium (Gibco #12200036) supplemented with 10% heat-inactivated fetal bovine serum and 100 U/mL penicillin, 100 ug/mL streptomycin. K562 were either subcultured at a 1:10 or 1:20 ratio or resuspended at 1E5 cells/mL every 2-3 d. NK92 (ATCC CRL-2407) were cultured in MEM α (Gibco 12000-022) with 0.2 mM Myo-inositol (Sigma I-7508), 0.1 mM 2-mercaptoethanol (Gibco 21985-023), 0.02 mM folic acid (Sigma F-8758), 1.5g/L sodium bicarbonate, 12.5% fetal bovine serum (Gibco 16000044), 12.5% horse serum (Gibco 16050122), and 100 U/mL penicillin, 100 µg/mL streptomycin. To culture, cells were kept in T25 or T75 flasks (Corning 431464U, 431463) and maintained between a density of 2×10^5^ cells/mL – 1×10^6^ cells/mL. Media was supplemented with 100 IU/mL recombinant IL2 (Peprotech 200-02) for parental (unmodified), mbSC BB, sSC BB, sp^null^3xFLAG-Neo2/15, eIL12, sp^null^3xFLAG-DR18, and sp^IgE^3xFLAG-DR18 NK92.

### Calcium phosphate transfection

Plasmid DNA in nuclease free water was mixed with 2 M CaCl_2_ (final concentration 0.3 M), pipetted 8 times to mix, then added dropwise to an equal volume of 2X HEPES buffered saline (280 mM NaCl, 50 mM HEPES, 1.5 mM Na_2_HPO_4_, pH 7.1) and pipetted up and down 4 times to mix. After 3 min and 30 s, the mixture was pipetted vigorously 8 times, then added dropwise to cells. Cell medium was changed to fresh medium after 8-16 h.

### Synthetic cytokine conditioned medium production

4.5×10^6^ HEK293FTs were plated in 10 cm dishes and allowed to adhere for 8 h. Cells were transfected with 20 µg of cytokine plasmid DNA and 1 µg of a plasmid encoding dsRedExpress2 using the calcium phosphate method. 24 h after medium change, supernatant was removed, spun 125xg 5 min at 4 C, then filtered through a 0.45-micron filter. Medium was used within a week of harvest.

### NK92 cell line generation

5×10^6^ HEK293FTs were allowed to adhere for 8 h in 10 cm dishes, then transfected with 8 µg psPAX2 (Addgene 12260), 3 µg PMD2.G (Addgene 12259), 10 µg transfer plasmid, and 500 ng plasmid encoding dsRedExpress2^68^ as a transfection control. Two 10 cm dishes were transfected per lentivirus type. The following morning, medium was changed to 10 mL fresh DMEM and plates were incubated for 28 h. Lentivirus-containing supernatant was pulled up from plates, centrifuged at 500xg for 2 min at 4 °C, and passed through a 0.45-micron filter. 4 mL of NK92 (1×10^6^ cells per mL in complete MEM α) was added to the lentiviral supernatant, along with 4 µg per mL of polybrene in 0.9% saline (EMD Millipore TR-1003-G). Cells were spun in viral supernatant at 37 °C for 90 min at 2500xg in a fixed angle rotor, then resuspended in viral supernatant and incubated overnight. The following morning, cells were spun at 125xg for 5 min at 4 °C, viral supernatant was aspirated, and cells resuspended in fresh media. Transduction was analyzed 2 d later (BD LSRFortessa SORP Cell Analyzer). To select for cells that expressed sSCs, cells were cultured with puromycin (1 µg/mL), then enriched by fluorescence-activated cell sorting (BD FACSAria IIu Cell Sorter). NK92 transduced with the mbSC BB were selected with blasticidin (10 µg/mL) for 14 days. NK92 transduced with mbIL2 were selected via culture without IL2 followed by selection with blasticidin (10 µg/mL) for 14 days. NK92 transduced with mbIL15 were selected via culture without IL2.

### K562 transduction

5×10^6^ Lenti-X HEK293Ts were plated in 10 cm dishes and allowed to adhere for 24 h. 8 µg psPAX2 (Addgene 12260), 3 µg PMD2.G (Addgene 12259), 10 µg transfer plasmid, and 1 µg plasmid encoding dsRedExpress2^68^ as a transfection control. The following morning, media were replaced with fresh medium. Cells were incubated for 32 h. Lentivirus-containing supernatant was collected from plates, centrifuged at 500xg for 2 min at 4 °C, and passed through a 0.45-micron filter. 1 mL of virus was added to 100 µL of K562 (1×10^6^ cells/mL) in a 12 well plate. Transduction was analyzed 3 d later (BD LSRFortessa SORP Cell Analyzer).

### Western blot analysis of protein expression

For analysis of EV markers, NK92 were seeded at 4×10^5^ cells/mL in incomplete MEM α and after 2 d, culture was centrifuged at 125xg 5 min to separate NK92 cells and conditioned medium. Vesicles were isolated from conditioned medium as described below but, after size exclusion chromatography, were concentrated with filters that had not been treated with bovine serum albumin. NK92 cells were rinsed with PBS and incubated with radioimmunoprecipitation assay buffer (RIPA buffer; 150 mM NaCl, 50 mM Tris-HCl pH 8.0, 1% Triton X-100, 05% sodium deoxycholate, 0.1% SDS) with protease inhibitor cocktail tablet (Pierce PIA32953; 1 tablet per 10 mL RIPA buffer) for 5 min at room temperature (∼20–25° C), incubated on ice for 30 min, then spun at 12,200xg for 20 min to clear supernatant, which was used in subsequent analyses. To produce cell lysates for synthetic cytokine analysis, 4.5×10^6^ HEK293FTs were transiently transfected with 20 µg of cytokine plasmid DNA using calcium phosphate transfection. 36 h after media change, cell lysate was harvested as done for NK92. Cell lysate protein content was analyzed using a bicinchoninic acid assay (Pierce). Lysate or vesicles were incubated with either reducing or non-reducing Laemmli buffer at either 70 or 90 °C for 10 min (see **Table 1**). For synthetic cytokine expression analysis, 5 µg of protein per well was loaded into a 4-15% polyacrylamide gradient Mini-PROTEAN TGX precast protein gel (Bio-Rad). For vesicle marker analysis, either 3 µg of protein from cell lysates or 4.8×10^8^ vesicles per well were loaded. Gels were run at 50 V for 10 min, then 100 V for 60 min at room temperature. Protein was then transferred to a PVDF membrane (Bio-Rad) at 100 V for 45 min. Membranes were blocked in either 3% non-fat milk in tris buffered saline (TBS; 50 mM Tris, 138 mM NaCl, 2.7 mM KCl, pH 8.0) for 30 m at room temperature (3xFLAG) or 5% non-fat milk in tris buffered saline plus tween (TBST; 50 mM Tris, 150 mM NaCl, 0.1% Tween 20, pH 7.6) for either 1 h at room temperature (Myc, Calnexin, CD81) or overnight at 4 °C (IL12p40, eIL15), then incubated with primary antibody (see **Table 1**). For 3xFLAG immunoblotting only, membranes were rinsed 3×5 min with TBS, with 0.05% Tween 20 added in the second and third rinse, prior to addition of primary antibody. After incubation with the primary antibody, all membranes were rinsed 3 x 5 min with TBST, then incubated with secondary antibody for 1 h room temperature. Afterwards, membranes were rinsed 3 x 5 min in TBST prior to addition of Clarity Western ECL reagent (BioRad) and subsequent exposure to film or chemiluminescence imaging with an Azure c280 imager.

**Table 1:** Antibody properties and sample preparation methods used in western blot analysis.

| Antibody target | Product no. | Laemmli buffer | Denature temperature (°C) | Antibody dilution | Animal of origin |
| --- | --- | --- | --- | --- | --- |
| 3xFLAG | Sigma F1804 | reducing | 70 | 1:1000 | mouse |
| Myc | Abcam ab32 | reducing | 70 | 1:1000 | mouse |
| IL12p40 | Abcam ab106270 | reducing | 70 | 1:1000 | rabbit |
| IL15 | Abcam ab7213 | reducing | 70 | 1:1000 | rabbit |
| Calnexin | Abcam ab22595 | reducing | 70 | 1:1000 | rabbit |
| CD81 | Santa Cruz sc23962 | non-reducing | 90 | 1:500 | mouse |
| Mouse | Cell Signaling 7076 | N/A | N/A | 1:3000 | horse |
| Rabbit | Invitrogen 32460 | N/A | N/A | 1:3000 | goat |

### NK92 irradiation

NK92 resuspended in MEM α were irradiated with 10 Gy of X-ray irradiation at a rate of 208 cGy/min, using a RadSource RS-2000 irradiator set to 160 kV and 25 mA, then immediately used in assays.

### Flow cytometry-based cytotoxicity assays

NK92 were incubated without IL2 for 1 d, then resuspended in MEM α at the maximum effector concentration tested, and serially diluted 2-fold to make cell suspensions for lower effector concentrations. 100 μL of NK92 in MEM α, or MEM α alone, were added to 100 μL of mScarlet-I^69^ expressing K562 resuspended in IMDM at 1×10^5^ cells/mL, in a 96 well plate. Cells were incubated for 20 h at 37°C, 5% CO_2_, and either ambient oxygen (∼21% O_2_) or hypoxia (1% O_2_), using a hypoxic incubator (HERAcell150i, Thermo Fisher). 3 μM DAPI and PKH26 reference microbeads (sigma P7458) were added to each sample, and samples were analyzed using a BD LSRFortessa SORP Cell Analyzer (**Supplemental Fig. 2A**). Live K562 cells (DAPI^neg^, mScarlet-I^pos^) counts were analyzed and normalized to the number of reference microbeads counted, to obtain a count of live tumor cells per well. For cytotoxicity assays, percent growth inhibition was defined as follows:

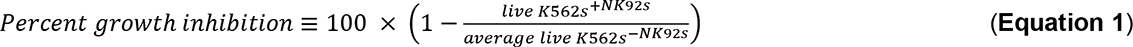

### Flow cytometry-based growth assays

Unirradiated NK92, or NK92 irradiated as described above, were resuspended at 2×10^5^ cells/mL. Cells were plated in two sets of 96 well plates (200 μL per well), then incubated at 37°C, 5% CO_2_ and either ambient O_2_ or 1% O_2_. After 2 d or 4 d, 3 μM DAPI and sigma PKH26 reference microbeads (sigma P7458) were added to one set of plates, and each sample was analyzed using a BD LSRFortessa SORP Cell Analyzer (**Supplemental Fig. 2A**). The number of live NK92 detected was normalized to the number of reference microbeads detected to obtain live NK92 counts per well. For consortia cell expansion assays, normalized cell count was calculated with the following formula:

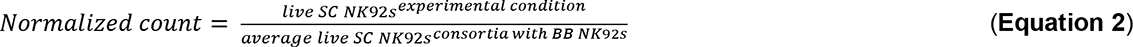

### Time lapse microscopy killing assays

NK92 were incubated for 1 d without IL2. K562 transduced to express mScarlet-I^69^ were resuspended in IMDM at 1×10^5^ cells/mL. For 4 h videos of killing, 3 μM DAPI was added to each well and wells were imaged every 5 min for 4 h total with a 10x objective, an ET-dsRed filter (Chroma 49005), a DAPI filter (Keyence OP-87762), and either a Cy5.5 filter (Chroma 49022) to detect mbSC NK92 or an EYFP filter (Chroma 49003) to detect sSC NK92. 100 μL of K562 per well were plated in a 96-well plate. NK92 were resuspended in MEM α at 3.125×10^5^ cells/mL and 100 μL of NK92 were added to K562, for a total volume of 200 μL. For 20 h killing kinetic assay, the plate was imaged with a 10x objective and an ET-dsRed filter every 2 h for 20 h total at five points per well in a time-lapse microscope (Keyence BZ-X800) fitted with an incubation chamber (Tokai Hit). Image-based cell-cytometry software (Keyence) was used to calculate tumor cell counts.

### Extracellular vesicle isolation

NK92 plated at 4×10^5^ cells/mL were cultured for 2 d in 10 mL of incomplete MEM α. Conditioned medium was centrifuged at 300xg for 10 min, and supernatant was further centrifuged at 2000xg for 20 min to remove dead cells and apoptotic bodies (Beckman Coulter Avanti J-26XP centrifuge, J-LITE JLA 16.25 rotor). Supernatant was centrifuged at 26,500 rpm for 2 hr 21 min (Beckman Coulter Optima L-80 XP ultracentrifuge, SW41 Ti rotor), using polypropylene ultracentrifuge tubes (Beckman Coulter 331372). All centrifugation steps were performed at 4 °C. Supernatant was aspirated until ∼100 μL remained and EV pellet was left on ice for 30 min. Resuspended EVs were transferred to microcentrifuge tubes, then run on a size exclusion chromatography column (Izon ICO-70), using PBS as a running buffer. The first 2 mL of eluent were collected and re-concentrated using 50 kDa ultrafilters (Amicon UFC8050) that been precoated with 1% bovine serum albumin in PBS for 1 h at room temperature; eluent was centrifuged for 30 min at 4000xg, 4°C to concentrate the EVs.

### Nanoparticle tracking analysis (NTA)

Vesicle concentration and size were measured using a Nanosight NS300 (Malvern) running software v3.4 and a 642 nm laser. Vesicles were diluted to 2-10 x 10^8^ particles/mL in phosphate buffered saline (PBS) for analysis. Samples were run at an injection rate of 30, imaged at a camera level of 14, and analyzed at a detection threshold of 7. Three 30 second videos were captured for each sample to determine the average vesicle concentration and size histograms.

### Transmission electron microscopy (TEM)

10 μL of purified EVs was placed onto a carbon-coated copper grid (Electron Microscopy Services, CF400-Cu-50) for 10 min before excess liquid was wicked away with a piece of filter paper. The grid was dipped in PBS twice to remove excess proteins and unreacted ligands from the media and reaction, and was allowed to dry for 2 min. To achieve negative staining, 10 μL of uranyl acetate solution (2 wt% in Milli-Q water) was placed on the grid for 1 min before being wicked away with filter paper. The grid was allowed to fully dry (3 h to overnight) at room temperature (approximately 20°C). Bright-field TEM imaging was performed on a JEOL 1230 TEM. The TEM operated at an acceleration voltage of 100 kV. All TEM images were recorded by a Hamamatsu ORCA side-mounted camera or a Gatan 831 bottom-mounted CCD camera, using AMT imaging software.

### Serial filtration of mbSC-conditioned media

Transfected HEK293FTs conditioned media for 28 h. Cells were removed from medium by centrifugation at 125 x g for 5 min, then filtration through a 0.45-micron filter. Conditioned media were serial filtered by first centrifuging at 4000 x g for 20 min in a 100 kDa molecular weight cutoff filter (Amicon UFC9100) then centrifuging at 4000 x g for 10 min in a 50 kDa filter (Amicon UFC9050) and finally, centrifuging at 4000 x g for 30 min for 10 kDa device (Amicon UFC9010). All centrifugation was performed at 4 °C.

## Supplementary Material Online

Online materials include Supplementary Information and Supplementary Videos (see **Supplementary Note 3** for descriptions). Source data will be made available with the final publication.

## Acknowledgements

This work was funded by in part by an AbbVie pilot project, a Lurie innovation award, and the National Institute of Biomedical Imaging and Bioengineering of the NIH under award number 1R01EB026510. This work was supported by the Northwestern University Center for Genetic Medicine and Robert H. Lurie Comprehensive Cancer Center Flow Cytometry Core Facility. The Lurie Cancer Center is supported in part by an NCI Cancer Center Support Grant #P30 CA060553.

## Authors Contributions

S.D. designed and performed experiments, as well as prepared the manuscript. P.S.D., J.A., and B.Z. contributed to experimental design. R.E.M. performed experiments. I.J.H. aided in execution of experiments. J.N.L. contributed to experimental design and manuscript preparation.

