## Supplementary Information for "Comparative evaluation of synthetic cytokines for enhancing production and performance of NK92 cell-based therapies"

**Contents:**

Supplemental Figures 1-3
Supplementary Notes 1-3

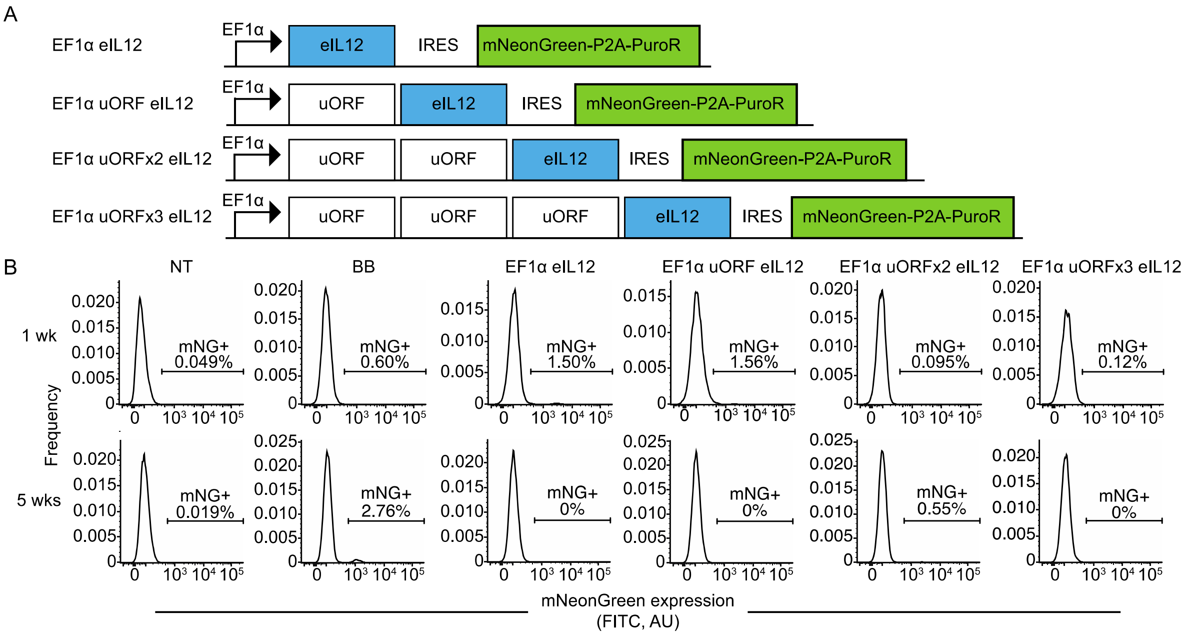


**Supplementary Figure 1: Upstream open reading frames enable long term NK92 expression of eIL12. A.** Schematic of eIL12 transgenes. 0, 1, 2, or 3 synthetic upstream open reading frames (uORFs) were inserted upstream of eIL12 and subsequent transduction markers to diminish transgenic protein expression to varying degrees, with the expectation that inclusion of more uORF sequences would confer greater reductions in translation initiation efficiency. Each uORF comprises the DNA sequence ATG-GGT-TGA. **B.** Persistence of eIL12 transduced NK92 after antibiotic selection. eIL12 or expression backbone-transduced NK92 were selected with puromycin, and transduction (mNG fluorescence) was assessed after 1 wk and 5 wks by flow cytometry. Gates were drawn to include < 0.05 % non-transduced cells, based on non-transduced NK92 at each time point. Gates are equivalent for panels within each time point. While at 1 wk of selection a small percentage of NK92 transduced with all eIL12 constructs could be detected, after 5 wks of selection only NK92 transduced with the expression backbone or EF1α uORFx2 eIL12 were detectable. Abbreviations: arbitrary units (AU), a transgene expression backbone identical to the EF1α eIL12 construct except that it lacks eIL12 (BB), elongation factor 1 α promoter (EF1α), encephalomycocarditis virus internal ribosome entry site (IRES), the green fluorescent protein mNeonGreen (mNG), non-transduced (NT), 2A self-cleaving peptide from porcine teschovirus-1 polyprotein (P2A), puromycin antibiotic resistance gene (PuroR), upstream open reading frame (uORF), week (wk).

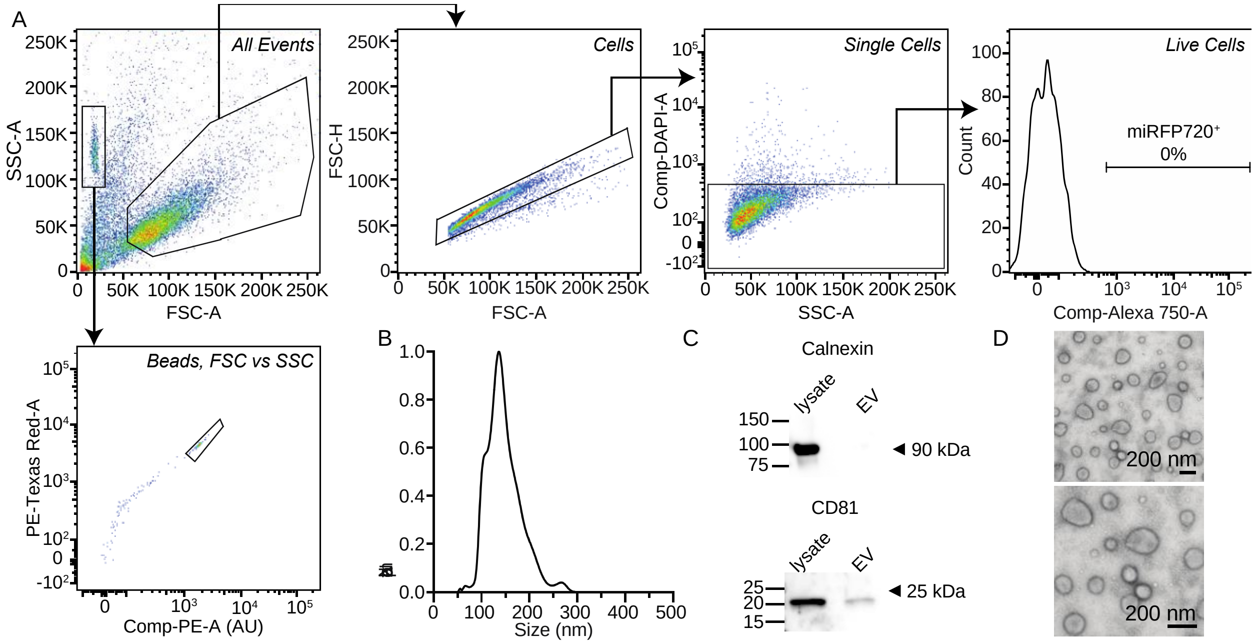


**Supplementary Figure 2: Methodological details—flow cytometry gating and validation of EV markers following reported purification method. A**. Representative strategy for gating live cells and PKH26 counting beads. Cells and counting beads were identified based on their FSC-A vs SSC-A profiles. Next, single cells were identified by further gating cells on FSC-A vs FSC-H. Next, live cells were identified by further gating single cells on SSC-A vs. DAPI-A. Live cells were then gated for fluorophore expression (e.g., miRFP720 expression), drawing gates such that <1% of fluorophore negative cells were included in the gate. PKH26 counting beads were better distinguished from debris by further gating counting beads on PE Texas Red-A vs Comp-PE-A. All fluorescent channels for fluorophores expressed by cells (DAPI-A, Alexa750-A, PE-A, FITC-A) were compensated as needed. **B.** Size distribution analysis of NK92-derived EVs. NK92-derived EVs were analyzed by NanoSight nanoparticle tracking analysis (NTA) and showed a mean size of 146.8 nm with a standard deviation of 35.6 nm, which is a size distribution typical for extracellular vesicles (as quantified by NTA, specifically). **C.** EV marker analysis. EVs were purified as described in **Methods**. NK92 cell lysates and NK92-derived EVs were analyzed with western blots against calnexin (a chaperone protein that is retained in the endoplasmic reticulum and is thus depleted in EV samples compared to whole-cell samples) and CD81 (a tetraspanin protein that traffics to EVs and is thus present in EV samples, as well as whole-cell samples). Arrows indicate expected sizes for each protein. EVs show expected pattern of calnexin and CD81 expression. **D.** Evaluation of NK92-derived EV morphology. NK92-derived EVs were imaged by transmission electron microscopy following negative staining (see **Methods**). EVs show a cup-like morphology typical of EVs evaluated by this method. Abbreviations: area (A), arbitrary units (AU), compensated (comp), extracellular vesicle (EV), forward scatter (FSC), height (H), kilodalton (kDa), nanometer (nm), side scatter (SSC).

**Supplementary Note 1. ANOVA results**

<u>Figure 1:</u>
Fig. 1H: One-way ANOVA: $p < 0.0001$
Fig. 1I: One-way ANOVA: $p < 0.0001$
Fig. 1J: One-way ANOVA: $p = 0.0003$
Fig. 1K: Two-way ANOVA:
Interaction: 5.117% total variation, $p < 0.0001$
Row factor (IL2 dose): 45.42% total variation, $p <0.0001$
Column factor (cell line): 42.85% total variation, $p <0.0001$

<u>Figure 3:</u>
Fig. 3D: One-way ANOVA: $p<0.0001$

<u>Figure 5:</u>
Fig. 5E: 3xFLAG-Neo2/15: One-way ANOVA: $p = 0.0167$
eIL12: One-way ANOVA: $p = 0.0009$
eIL15: One-way ANOVA: $p = 0.0046$
3xFLAG-DR18: One-way ANOVA: $p = 0.0005$

<u>Figure 6:</u>
Fig. 6B: mbSC no contact: One-way ANOVA: $p < 0.0001$
3xFLAG-Neo2/15, no contact: One-way ANOVA: $p < 0.0001$
eIL15, no contact: One-way ANOVA: $p < 0.0001$
Fig. 6D: One-way ANOVA: $p = 0.464$

<u>Figure 7:</u>
Fig. 7D: One-way ANOVA: $p < 0.0001$
Fig. 7E: Two-way ANOVA:
Interaction: 18.31% total variation, $p < 0.0001$
Row Factor: 33.28% total variation, $p < 0.0001$
Column Factor: 47.37% total variation, $p < 0.0001$

<u>Figure 8:</u>
Fig. 8A: Two-way ANOVA:
Interaction: 5.552% total variation, $p = 0.6858$
Row Factor (sSC line): 48.16% total variation, $p = 0.0006$
Column Factor (mbSC line): 6.746% total variation, $p = 0.0374$
Fig. 8B: Two-way ANOVA:
Interaction: 22.50%, $p < 0.0001$
Row Factor (sSC line): 43.40, $p < 0.0001$
Column Factor (mbSC line): 19.42, $p < 0.0001$
Fig. 8C: Two-way ANOVA:
Interaction: 18.25%, $p < 0.0001$
Row Factor (sSC line): 9.018%, $p < 0.0001$
Column Factor (consortia condition): 69.46%, $p < 0.0001$

**Supplementary Note 2. Time scale of sSC NK92 sorting**

This note provides additional details for the experiments presented in **Figure 2F**. sSC NK92 were sorted at various time points prior to analysis. sSC BB and eIL15 NK92 were thawed from existing cell stocks that had been sorted in prior work. sp$^{null}$3xFLAG-DR18 NK92 were sorted 23 d before day 0. All other sSC NK92 lines were sorted 10 d prior to day 0.

**Supplementary Note 3. Supplementary Video descriptions**

Supplementary Video 1: NK92 growth: mbSC backbone

Supplementary Video 2: NK92 growth: mbIL2

Supplementary Video 3: NK92 growth: mbIL15

Supplementary Video 4: killing assay: K562 only

Supplementary Video 5: killing assay: K562 + mbSC backbone NK92

Supplementary Video 6: killing assay: K562 + mbIL2 NK92

Supplementary Video 7: killing assay: K562 + mbIL15 NK92

Supplementary Video 8: killing assay: K562 + sSC backbone NK92

Supplementary Video 9: killing assay: K562 + 3xFlagNeo2-15 NK92

Supplementary Video 10: killing assay: K562 + IgEsp3xFlagNeo2-15 NK92

Supplementary Video 11: killing assay: K562 + eIL12 NK92

Supplementary Video 12: killing assay: K562 + 3xFlagDR18 NK92

Supplementary Video 13: killing assay: K562 + IgEsp3xFlagDR18 NK92

Supplementary Video 14: killing assay: K562 + eIL15 NK92
